# Overexpression of ADAR1 in mice does not initiate or accelerate cancer formation *in vivo*

**DOI:** 10.1101/2023.01.29.526130

**Authors:** Shannon Mendez Ruiz, Alistair M Chalk, Ankita Goradia, Jacki Heraud-Farlow, Carl R Walkley

**Author notes:** Joint authors (Jacki Heraud-Farlow, Carl R Walkley).

## Abstract

Adenosine to inosine editing (A-to-I) in regions of double stranded RNA (dsRNA) is mediated by adenosine deaminase acting on RNA 1 (ADAR1) or ADAR2. ADAR1 and A-to-I editing levels are increased in many human cancers. It is not established if elevated ADAR1 represents a driver or passenger during cancer formation. We established a series of murine alleles to allow *in vivo* overexpression of ADAR1, its individual isoforms or mutant forms of ADAR1 to understand how it contributes to cancer pathogenesis. The widespread overexpression of ADAR1 or either the p110 or p150 isoforms as sole lesions was well tolerated and did not result in cancer formation. Therefore, ADAR1 overexpression alone is not sufficient to initiate cancer. We demonstrate that endogenous ADAR1 and A-to-I editing levels increased upon immortalization by loss of p53 in murine cells, consistent with the observations from human cancers. We tested if ADAR1 overexpression could co- operate with cancer initiated by loss of tumour suppressors using a model of osteosarcoma. We did not see a disease potentiating or modifying effect of overexpressing ADAR1 or its isoforms. We conclude that the increase in ADAR1 expression and A-to-I editing in cancers is a passenger, rather than a driver, of tumor formation.

## INTRODUCTION

It is now appreciated that RNA undergoes extensive post-transcriptional modification, collectively referred to as the RNA epitranscriptome (1, 2). These modifications can be transient and written/erased or can result in a permanent change to the RNA. These changes can influence the protein coding potential of the transcript, as well as other critical aspects of the RNA lifecycle including localisation, structure, interactions with other RNA binding proteins/RNAs and phase separation (3). An increasing body of work demonstrates that the RNA of cancer cells, like normal cells, undergoes extensive modification (4–7). Detailed studies of the epitranscriptome of cancer has determined that there are changes in RNA modifications between cancer cells and normal tissue. In parallel, forward genetic studies have determined that enzymes catalysing the modifications of RNA may be unique vulnerabilities in cancer cells (8–12). This has led to a high interest in the therapeutic modulation of RNA modifying enzymes for a range of cancer types and applications (13, 14).

One of the most prevalent epitranscriptomic modifications in mammals is the conversion of adenosine to inosine (A-to-I editing) in regions of double stranded RNA (dsRNA) (15–18). A-to-I editing results in the permanent change of the RNA sequence, with no currently defined reversion enzymes/mechanisms known. There are millions of A-to-I editing events in the human transcriptome, with editing occurring across the length of an RNA (19–23). *In vivo* the levels of editing at any given adenosine can vary from very low to 100% of the RNA molecules, however, the majority of sites are lowly edited (19, 21). A-to-I editing occurs in regions of dsRNA structure and in mammals is catalysed by Adenosine Deaminase Acting on RNA 1 (ADAR1) and ADAR2 (17). A well characterised consequence of A-to-I editing by ADARs is a change in the protein coding sequence, yielding a different protein or a protein with altered function from the genomic encoded sequence. This “recoding” activity of ADARs has been demonstrated for both ADAR1 and ADAR2 and can have profound physiological impact (24). Editing can also occur at multiple sites within a short stretch of a dsRNA sequence (termed “hyperediting” or “repetitive region editing”), which is mostly associated with editing of repetitive elements such as *Alu* (primate restricted) and the related B1/B2 SINEs in rodents (19,22,25–28). Most editing in mammals occurs in repeat sequences.

Despite the diverse range of potential functions and consequences of A-to-I editing it has now been established that ADAR1 is an essential regulator at the interface of cellular dsRNA and the innate immune sensing system. The major physiological function of ADAR1 is to edit the cells own dsRNA to prevent them being sensed as “non-self” by the cytoplasmic innate immune sensor MDA5 (29–32). This has been established in mouse and human (27,33,34). Genetic analysis in the mouse, and more recently in human cell lines, has further refined the understanding of the nexus between ADAR1 and the innate immune system to demonstrate the critical role for the cytoplasmic ADAR1p150 isoform (35–37). These studies have demonstrated that the loss of ADAR1p150, but not the nuclear ADAR1p110 isoform, leads to activation of the innate immune response to cellular dsRNA ligands. Whilst the physiological significance of editing by ADAR1 of cellular dsRNA has been established, it is not yet determined how this activity contributes in pathological states such as cancers.

Changes in A-to-I editing were recognized in cancer transcriptomes prior to the availability of current sequencing methods (38). Initial reports of A-to-I editing in cancer described changes, generally reductions, of ADAR2 mediated editing at selected targets in tumours of the CNS such as glioblastoma and astrocytoma (39). Recent studies utilising large RNA-seq datasets from diverse human cancers have identified a trend of increased overall editing and ADAR1 expression in cancer types ranging from leukaemia to solid tumors (40–45). Single cell analysis of lung cancer confirmed that there was cancer cell intrinsic elevated A-to-I RNA editing (46). These studies have identified a correlation between increased ADAR1 expression and increased editing in cancer. Additional studies have extended this analysis to demonstrate changes in cancer proteomes arising from A-to-I editing (47). Increased ADAR1 expression has been associated with either copy number gains at Chromosome 1, where the *ADAR* gene is located in humans, or the activation of interferon/innate immune sensing responses in tumors leading to an increase in ADAR1 expression from the IFN-inducible promotor upstream of ADAR1p150 (42). The biological consequences of increased ADAR1 and an increased level of overall editing in tumors is only beginning to be explored. In some specific examples, such as in melanoma, reduced editing efficiency has been proposed to be important in the pathogenesis of these tumours (48), although this appears to be less common than an increased expression of ADAR1 and higher overall A-to-I editing levels. In other examples it has been proposed that protein recoding of specific RNAs is a key function of ADAR1 in evolution and prognosis of gastrointestinal tumours or hepatocellular carcinoma (41, 49). More recently, the genetic loss of ADAR1 was demonstrated to be highly sensitizing to immune checkpoint blockade in *in vivo* genome-wide screens (8,10,12). This has now been replicated across a broader range of tumor types (50–52) and in extended contexts (53), suggesting inhibition or loss of ADAR1 may be a broadly applicable therapeutic cancer strategy. These studies have demonstrated that targeting ADAR1 may yield direct cell intrinsic benefits in addition to boosting immune response against tumours. ADAR1 loss has also been implicated in enhancing the response to therapies such as demethylating agents (54). We are only just beginning to understand the consequences of changes in A-to-I editing on cancer initiation and maintenance, both at the level of its effect on specific transcripts and also on the global transcriptome of the cancer cells. How ADAR1 contributes to tumor initiation and evolution requires further study (Fig 1A).

**Figure 1.**
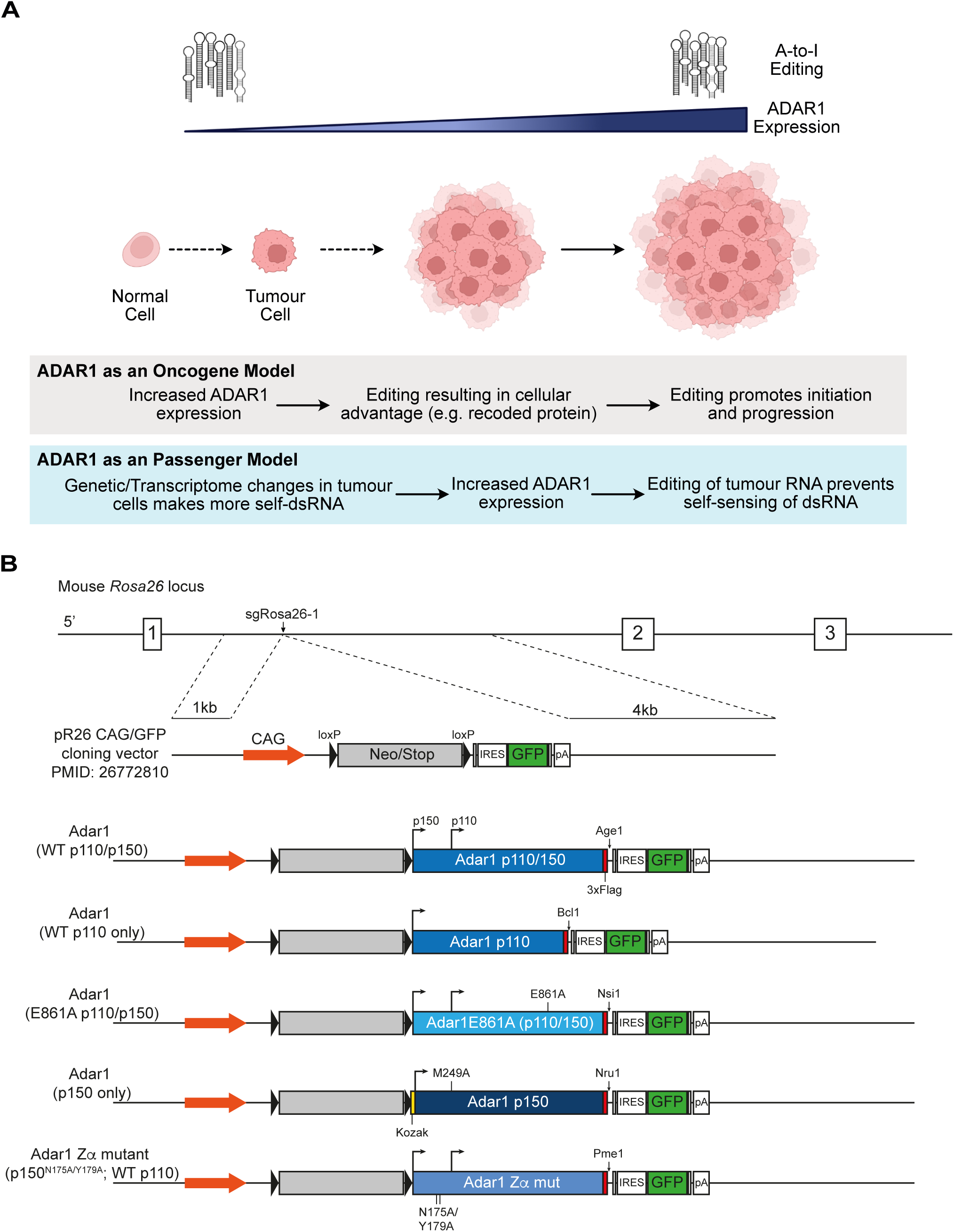
Proposed model of ADAR1 action in cancer. **(A)** Proposed models of the action of ADAR1 in cancer. In the “ADAR1 as an oncogene” model, elevated ADAR1 leads to increased A-to-I editing and this acts as a tumor initiating event or promotes tumor establishment and maintenance. In the alternate “ADAR1 as a passenger model” ADAR1 is elevated as a result of changes in the tumor transcriptome and environment, leading to increased ADAR1 as a secondary consequence. **(B)** Schematic of the constructs used to overexpress murine *Adar1* cDNA from the Rosa26 locus in mice.

Here we report the experimental evaluation of the role and effects of over-expression of ADAR1 *in vivo* on cancer initiation and progression. We generated a series of knock-in mouse models where we can ectopically overexpress ADAR1, its specific isoforms or mutant forms, *in vivo*. All cDNA are expressed from the same locus allowing us to determine if over-expression of ADAR1 *in vivo* is able to act as a tumor initiating event/founder lesion. We also assessed if increasing ADAR1 levels can co-operate by potentiating tumor development or metastatic spread in a murine model of human solid cancer. From these studies we conclude that over-expression of ADAR1 as a sole lesion is not sufficient to initiate cancer *in vivo*. We find that the increased expression of ADAR1 or its individual isoforms is well tolerated long term without any increase in cancer incidence. We further observed that the over-expression of ADAR1 did not modify the frequency of primary tumors or metastatic frequency in a model of osteosarcoma driven by the loss of *Trp53* and *Rb1* (55). We conclude that overexpression of ADAR1 does not initiate cancer or co-operate to promote tumor formation *in vivo*. Our analysis indicates that increased ADAR1 in cancer is a consequence of cancer formation and the modified cancer dsRNA transcriptome.

## MATERIAL AND METHODS

### Ethics Statement

All animal experiments conducted for this study were approved by the Animal Ethics Committee of St. Vincent’s Hospital, Melbourne, Australia (Protocol number #009/18 and #030/21). Animal studies were conducted in accordance with the Australian code for the care and use of animals for scientific purposes, 8th Edition (2013; ISBN:1864965975). Animals were euthanized by cervical dislocation or CO_2_ asphyxiation.

### Biological Resources

#### Murine *Adar1* expression constructs

Murine *Adar1* cDNA was generated by gene synthesis (GeneArt, Germany) based on the CCDS sequence for murine full length ADAR1 (encoding p110 and p150) from the C57Bl/6 reference strain. A C-terminal 3xFlag sequence added to all constructs by subcloning of gBlocks (IDTDna, Singapore). All mutations were introduced by subcloning of gBlocks and all final constructs were sequence verified. The following constructs were made: full length *Adar1* (expressed both p150 and p110); p110 only expressing; p150 only expressing construct had a consensus Kozak sequence (AGCCACC) inserted immediately prior to the initiation codon in place of the native murine sequence and a M249A mutation to disrupt the p110 initiation codon (56); editing deficient has an E861A mutation (29); Za mutant had compound N175A/Y179A mutations introduced as previously described (53). All constructs were cloned into pLVX-IRES-Puro (modified from pLVX vector (Clontech) by removal of the ZsGreen and replacement with a multiple cloning site and IRES-Puro; provided by S Mannering, St Vincent’s Institute). These constructs were packed into lentivirus by transient transfection in 293T cells with psPAX2 (Addgene) and the ecotropic MLV envelope pCAG-Eco using calcium phosphate as described (57). psPAX2 was a gift from Didier Trono (Addgene plasmid # 12260; http://n2t.net/addgene:12260; RRID:Addgene_12260) and pCAG-Eco was a gift from Arthur Nienhuis and Patrick Salmon (Addgene plasmid # 35617; http://n2t.net/addgene:35617; RRID:Addgene_35617). Virus containing supernatant was collected, centrifuged and stored at -80C until required.

To confirm expression, the lentivirus was used to infect the murine stromal cell line Kusa4b10 (58, 59). Cells were spin infected in 6 well plates, expanded on 10 cm plates and treated with puromycin to select for infected cells. Once control cells had died, puromycin was removed and cells expanded and used for western blot analysis as described below.

#### Animals

The murine *Adar1* cDNA were cloned from the pLVX-IRES-Puro cloned into the pR26-CAG/GFP-Asc vector (60). pR26 CAG/GFP Asc was a gift from Ralf Kuehn (Addgene plasmid # 74285; http://n2t.net/addgene:74285; RRID:Addgene_74285). All targeting constructs also included an Adar1 cDNA specific unique restriction site between the *Adar1* cDNA and IRES-GFP (see Figure 1). This could be used to confirm the cDNA inserted if required on genomic DNA PCR products. *Rosa26*-Lox-Stop-Lox-*Adar1* mice were generated using CRISPR/Cas9 targeting in C57BL/6 zygotes by The Melbourne Advanced Genome Editing Center (MAGEC) at the Walter and Eliza Hall Institute (Parkville, Australia). All constructs were confirmed by Sanger sequencing and then after successful targeting by long range PCR, unique restriction digest and Sanger sequencing of the region in both the founders and subsequent generations. This established the five strains: *R26*-*Adar1^ki/+^* (expressed p110 and p150), *R26*-*Adar1p150^ki/+^* (p150 only), *R26*-*Adar1p110^ki/+^* (p110 only), *R26*-*Adar1E861A^ki/+^* (editing dead p110 and p150), *R26*-*Adar1-Zα^ki/+^* (N175A/Y179A p150 mutant; wild-type p110).

The *Ubiquitin C* (*Ubc*) promoter driven CreER^T2^ (*Ubc*-CreER^T2/+^) (61) was obtained from P Humbert (La Trobe University, Victoria) from stock originally purchased from The Jackson Laboratory (B6.Cg- *Ndor1^Tg(UBC-cre/ERT2)1Ejb^*/2J; stock#008085). We then cross-bred these mouse strains to create five Cre- inducible ADAR1 isoform mouse lines: *Ubc*-CreER^T2Tg/+^ *R26*-*Adar1^ki/+^*, *Ubc*-CreER^T2Tg/+^ *R26*- *Adar1p150^ki/+^*, *Ubc*-CreER^T2Tg/+^ *R26*-*Adar1p110^ki/+^*, *Ubc*-CreER^T2Tg/+^ *R26*-*Adar1E861A^ki/+^*, *Ubc*- CreER^T2Tg/+^ *R26*-*Adar1-Zα^ki/+^*. Wherever possible *Ubc*-CreER^T2Tg/+^ *R26*-wildtype littermate controls were housed together with experimental mice, administered tamoxifen containing food and used as age matched controls. For acute somatic deletion model (*Ubc*-CreER^T2^), all animals were ≥8 weeks of age at tamoxifen initiation. Tamoxifen containing food was prepared at 400mg/kg tamoxifen citrate (Selleckchem) in standard mouse chow (Specialty Feeds, Western Australia). For the osteosarcoma mouse model, *R26*-*Adar1^KI/+^* alleles were interbred with an established *Osx1*-Cre *Trp53*^fl/fl^ *Rb1^fl/fl^* mouse line that has been previously described (55,62,63). All mice were on a C57BL/6 background. All animals were housed at the BioResources Centre (BRC) at St. Vincent’s Hospital, Melbourne. Mice were maintained and bred under specific pathogen-free conditions with food and water provided *ad libitum*.

#### Genotyping

Tissue samples were collected into 1.5mL microcentrifuge tubes and spun at 17,000g for 2 minutes. They were then suspended in 300μL 50mM NaOH and digested at 95°C and 120rpm on a Vortemp56 Shaking incubator for 20 minutes. 100μL of 1M Tris-HCL (pH8.0) was added to the digested solution and vortexed for 5 seconds. The samples were spun again at 17,000g for 3 minutes on a Heraeus Multifuge 3SR+ centrifuge. At this stage the gDNA solution was used in the genotyping protocol described below. For adult mice, gDNA was extracted from ear clippings or other collected tissues with the DNeasy Blood and Tissue Kit (Qiagen), as outlined by the manufacturer.

For each reaction 1μL of gDNA solution was suspended in a PCR tube with 2μL of MyTaq red reaction buffer (5x; Bioline), 1µL of pooled primers/oligonucleotides (final concentration of each primer 10μM or 5μM for *Ubc*-Cre), 0.1μL of MyTaq HS DNA polymerase, and 5.9μL of nuclease-free water. All reactions were run on a Mastercycler Pro PCR machine (Eppendorf). 3% agarose gels were prepared with a 1:20000 dilution of SYBRSafe DNA gel stain (Invitrogen). Once complete the PCR products were loaded onto the gel and the electrophoresis was run on a Powerpac Electrophoresis system (BioRad) at 120 V for 45 minutes. Gels were then photographed using a VersaDoc (BioRad).

Sanger sequencing (Australian Genome Research Facility (AGRF), Melbourne) of the purified PCR product was performed as required. Recombination of the Lox-Stop/Neo-Lox cassette within the tissues of tamoxifen treated mice for Cre activated constructs was confirmed through PCR analysis. Recombinant targeting 10μM primer mixes were run with gDNA samples on the same PCR cycle specified in Supplemental Table 1. The *Ubc*-CreER^tg/+^, R26-*Adar1^ki/+^* constructs were amplified with a 10μM mix containing CAG For1, Neo Rev1, and one of either Adar1Rev1 (for full length, p150, E861A and Z*α* constructs) or Adar1p110 Rev1 for p110 overexpression mouse lineages. For p53 excision, a 5μM mix containing P53F2-10F and P53F2-10R targeting the Trp53fl/fl allele was compared against a 5μM mix of P53F2-10F, P53F2-10R, and P53F2-1F to target the excised allele. Genotyping of *Osx1*- Cre *Trp53*^fl/fl^ *Rb1^fl/fl^* was performed as previously described (55). Primer sequences in Supplemental Table 1.

#### Peripheral blood analysis

Peripheral blood (approximately 100μl) was obtained via retro-orbital bleeding. The blood was red blood cell-depleted using hypotonic lysis buffer (150mM NH_4_Cl, 10mM KHCO_3_, 0.1mM Na_2_EDTA, pH7.3) and resuspended in 50μl of FACS buffer for flow cytometry analysis.

#### Flow cytometry analysis

Bone marrow was harvested from both femurs of all collected mice. The femurs were flushed twice using a 1mL syringe and 23-gauge needle with 2mL of FACS buffer (PBS with 2%FCS). Spleen and thymus were removed, cleaned of connective tissue, weighed, and stored in FACS buffer. The tissues were then crushed against a 40μm cell strainer (Falcon, BD Bioscience) with the back of a 3mL syringe plunger in a 6-well plate containing either 5mL of FACS buffer for spleen, or 2mL per thymus. The flushed cell suspension was then filtered through 40μm cell strainers, with 100μL aliquoted into a 1.5mL microcentrifuge tube for counting on a Sysmex KX21 haematological analyser.

Single cell suspensions from peripheral blood, bone marrow, spleen and thymus were stained with antibodies (eBioscience, BioLegend or BD Pharmingen (29,59,64,65)) for flow cytometry (all antibody and conjugates in Supplemental Table 2). Cells were aspirated on a BD LSR II Fortessa and Cell Diva software version 8.1 (BD Biosciences, San Jose, CA, USA) was used for data acquisition and adjusting compensation. FlowJo software version 10.6.1 (Treestar) was used for sample analysis.

#### RNA-seq

Whole liver from tamoxifen treated mice of the indicated genotypes was used to purify RNA for RNA-seq. RNA samples were diluted in nuclease-free water to a concentration ranging between 100- 270ng/μL. The total RNA used for poly(A) enrichment and library preparation following by sequencing on the Illumina platform (150bpPE; library construction and sequencing performed by Novogene (Singapore)).

#### Gene Expression and Editing analysis

Pre-processing: Sequenced reads (150bp) were trimmed for adaptor sequence and low quality reads using fastp (v 0.19.5) (66). Parameters: --trim_front1 10 --trim_front2 10. Reads mapping to rRNA were removed using Bbmap (parameters: bbsplit.sh minratio=0.56 minhits=1 maxindel=16000) (67).

Differential expression: Counts were determined by mapping trimmed reads using Salmon (v1.4.0) vs mm10 gencode vm22 (68). Salmon count data was processed using tximeta (69), and summarised to gene using lengthScaledTPM. Normalisation, QC and filtering were performed in Degust (70). Genes were filtered (count > 1, min CPM > 1 in at least 2 samples) and differential expression was performed using voom with sample weights (71). Each comparison for the ER model (*Adar1^ki/+^* vs WT, *Adar1^p110ki/+^* vs WT, *Adar1^p150ki/+^* vs WT and *Adar1^E861Aki/+^* WT), and the OS model (D21 vs D7) was performed separately. See Supplementary Dataset 1,3.

Heatmaps: Tidyheatmap (72) was used to visualise datasets.

Gene sets: The interferon stimulated gene set was derived from Liu et al., (12).

Editing analysis: Mapping: Preprocessed reads were aligned to the MM10/GRCm38 reference genome with transcript annotation (gencode.mm10.vM14.annotation.SEQINS.gtf) with STAR (version 2.6.0c)(73) using the following parameters: --outFilterType BySJout --outSAMattributes NH HI AS NM MD --outFilterMultimapNmax 20 --outFilterMismatchNmax 999 --outFilterMismatchNoverReadLmax 0.04 --alignIntronMin 20 --alignIntronMax 1000000 --alignMatesGapMax 1000000 -- alignSJoverhangMin 8 --alignSJDBoverhangMin 1 --sjdbScore 1 --sjdbOverhang 149. Duplicate reads were marked with Picard [“Picard Toolkit.” 2019. Broad Institute, GitHub Repository. http://broadinstitute.github.io/picard/; Broad Institute].

Differential editing of known sites: A database of 135,697 murine editing sites was compiled from published databases (RADAR; (74)), publications (26,29,65) and unpublished murine datasets (JH-F, AMC and CRW) and the datasets assessed for editing at these sites. Calling of differential editing in known sites across genotypes was performed using JACUSA 2.0.0-RC5 (75). Briefly, call-2 was used to determine the difference in editing level for all known sites (all replicates of genotype A vs all replicates of genotype B). Duplicate reads were removed. Sites required ≥50 read coverage and an editing rate of ≥0.01 (≥1%) to be considered. See Supplementary Dataset 2, 4.

Annotation: Editing sites were annotated with gene, gene part (promoter, Exon, intron, 3’ UTR or intergenic) using Goldmine (76). B1 and B2 SINE annotation (mm10) was from UCSC rmsk table (77).

AEI: We calculated the editing of repetitive sequences using a murine modified version of the *Alu* editing index (AEI) (26,65,78).

#### Histology

As *Osx*-Cre *Trp53^fl/fl^ Rb1^fl/fl^* mice were euthanized and autopsied, tumour samples were either fixed for histology or snap frozen and stored at -80C. For histology, pieces of primary and metastatic tumours were fixed in 4% formalin PBS solution in 10mL falcon tubes for 24-72 hours and then transferred to 70% ethanol for storage. The fixed samples were processed (embedded, sectioned and H&E stained) by the O’Brien Institute histology lab (St Vincent’s Institute). Images of the sections were provided by J Palmer (Histology Laboratory Coordinator).

#### Protein extraction and Western blotting

Kusa4b10 cells were lysed in RIPA ice cold RIPA buffer lysate mix (20mM Tris·HCl, pH8.0, 150mM NaCl, 1mM EDTA, 1% sodium deoxycholate, 1% Triton X-100, 0.1% sodium deoxycholate, 1x Halt protease inhibitor; 1x PhosSTOP phosphatase inhibitor). For western blot analysis 15mg of protein was loaded onto the gel. Frozen tissue samples were transferred to a 10mL tube and suspended in 350μL of ice-cold RIPA buffer lysate mix. Keeping samples cold, the samples were homogenized on low-medium speed with a mechanical homogenizer (IKA T10 basic S5 Ultra-turrax Disperser) to disperse the tissue. The homogenizer was rinsed three times each between samples with H_2_O and 70% ethanol to prevent cross contamination between samples. Homogenised tissue samples were freeze-thawed for 5 minutes on dry ice and transferred onto ice. Thawed lysates were aliquoted into 1.5mL microcentrifuge tubes and spun at 13,000g for 20 minutes at 4°C. The supernatant was transferred into fresh 1.5mL microcentrifuge tubes and either stored at -80°C or set aside on ice for protein quantification. Frozen tissue pellets, such as the spleen and thymus, were resuspended in 100μL of the RIPA buffer mix and incubated on ice for 20 minutes before undergoing the freeze-thaw step and 20mg of protein lysate used.

Protein extracts were loaded on pre-cast NuPAGE™ 10%, Bis-Tris (1.5 mm, 10-well) polyacrylamide gels (Invitrogen) and transferred onto Immobilon-P PVDF membranes (Merck Millipore). Membranes were blocked with 5% milk in Tris-buffered saline with tween (TBST) and incubated at 4°C overnight with rat monoclonal anti-mouse ADAR1 antibody (1:500 on primary tissue westerns or 1:1000 dilution for cell lines; clone RD4B11; inhouse hybridoma purified by Monash Antibody Technology Facility; also available from Merck Millipore as MABE1790) (29), Monoclonal Anti-FLAG M2-Peroxidase (HRP) antibody produced in mouse (1:2000 dilution; Sigma Aldrich, A8592), mouse anti-Actin (1:5000 dilution; Sigma Aldrich, MS-1295-P0). Membranes were then probed with HRP-conjugated goat anti-rat (Thermo Fisher Scientific, 31470) or anti-mouse (Thermo Fisher Scientific, 31444) secondary antibodies and visualized using ECL Prime Reagent for chemiluminescent detection on Hyperfilm ECL (Amersham).

### Statistical Analyses

The statistical significance was determined through both one-way and two-way ANOVA, chi-squared test, JACUSA 2.0.0-RC5 statistic (likelihood ratio of two samples), (Kaplan-Meier survival plots on Prism software version 8 and 9 (GraphPad; San Diego, CA, USA). Z-tests were conducted through Microsoft Excel software version 16.53 (Microsoft, USA). Data is presented as mean ± Standard error of the mean (SEM). Sample populations are defined in their relevant figure legends. Significance is defined under the following conventions: *P<0.05, **P<0.01, ***P<0.001, ****P<0.0001.

### Data Availability/Sequence Data Resources

RNA-seq datasets generated for this study are available in NCBI GEO (Accession number: GSE221628).

## RESULTS

### Generation of conditional ADAR1 overexpressing mouse models

We propose that there are two models which could account for the observed elevation of A-to-I editing and ADAR1 in cancers that has been independently reported across diverse cancer types (Fig 1A). The first is that elevations in ADAR1 and its A-to-I editing activity act as a driver of tumor initiation and development (ADAR1 as an oncogene model). In this model, ADAR1 editing rewires the cellular transcriptome to either favor tumour initiation or allow progression when cancer initiation is driven by other lesions. The second model is that ADAR1 is elevated in response to transcriptional and environmental adaptions of the tumor and its elevation enables tumor immune evasion by editing of potentially immunogenic self-derived dsRNA that are unique to the tumor transcriptome (ADAR1 as a passenger model). The first model would predict that overexpression of ADAR1 would lead to acquisition of cancerous phenotypes or even be sufficient to initiate cancers *in vivo* as a single event. The second model would predict that overexpression of ADAR1 is not sufficient to initiate cancer and that it also would not strongly cooperate to allow cancer establishment and evolution. We sought to directly test these two models using *in vivo* overexpression of ADAR1 or its isoforms.

We first sought to establish murine cDNA constructs that could be used to specifically overexpress ADAR1 isoforms. To this end, we generated a murine cDNA for endogenous *Adar1* by gene synthesis. From this cDNA we generated a range of alternatives using short replacement synthetic sequences (gBlocks) that could be directly cloned into the parental construct. We additionally included a C-terminal 3xFlag tag to allow differentiation between the endogenous and the ectopic ADAR1 proteins. In addition to full length Adar1 (capable of expressing both p150 and p110 isoforms; FL), we generated a p110 isoform only expressing cDNA by removal of all sequence preceding the p110 initiation codon at M249 in the full-length murine sequence. We further generated an editing dead mutant (E861A) which renders both p150 and p110 isoforms editing deficient (29); a Z*α* mutant through introduction of compound N175A/Y179A mutations, the murine homologue of the human N173A/Y177A mutations which are known to ablate Z-form nucleic acid binding (79–81), as previously described (53). We also generated two different constructs that would be predicted to be only able to express the full length p150 isoform. This was done by introducing a consensus Kozak sequence (AGCCACC) immediately prior to the initiation codon in place of the native murine sequence, with a second cDNA containing both a consensus Kozak and a M249A substitution to mutate the p110 initiation codon (56). We tested these cDNA by viral mediated overexpression in the murine stromal cell line Kusa4b10. This demonstrated that all cDNA generated ADAR1 proteins of the expected size (Supplemental Fig 1). The p150 only expressing cDNA demonstrated that the provision of a consensus Kozak sequence is largely sufficient to prevent p110 expression from the full-length cDNA sequence consistent with that previously described (56). The M249A mutation in addition to the consensus Kozak was sufficient to abrogate p110 expression near completely and yield a p150 isoform only expressing construct (Supplemental Fig 1). This construct was used for all subsequent experiments.

We then cloned the FL, p110, E861A, Z*α* mutant and p150 (containing both a consensus Kozak and M249A) into the pR26-CAG/GFP-Asc vector (60) to allow targeting into the *Rosa26* locus. This vector has a loxP flanked stop cassette (lox-Stop-lox; LSL) followed by the *Adar1* cDNA then an internal ribosomal entry sequence (IRES) and green fluorescent protein (GFP; Fig 1B). All *Adar1* cDNA would be able to be expressed from the same locus *in vivo*, allowing a comparison between the effects of each *in vivo*. Furthermore, these would be able to be somatically expressed and as either a heterozygous or homozygous allele. We incorporated an *Adar1* construct unique restriction site between each *Adar1* cDNA and IRES-GFP to allow for allele specific restriction digest of genotyping PCR product if required (Fig 1B). Upon Cre mediated recombination of the Lox-Stop-Lox cassette, the cell would express the respective *Adar1* cDNA and GFP from the *Rosa26* locus. Using these constructs we generated knock-in alleles of each Adar1 construct at the *Rosa26* locus in C57BL/6 oocytes.

### *In vivo* overexpression of ADAR1

To enable widespread overexpression of ADAR1 we cross the R26-LSL-*Adar1* mice to the *Ubc*- CreER^T2^ allele (61, 82). This enabled the inducible overexpression of ADAR1 following tamoxifen treatment of the mice. We established cohorts of *Ubc*-CreER^T2^ R26 wild type animals (Cre+ve controls) and *Ubc*-CreER^T2^ R26-LSL-*Adar1* (Cre+ve ADAR1 overexpressing). We first isolated tail fibroblasts and treated the cells in vitro with tamoxifen and confirmed that all induced GFP in a tamoxifen dependent manner (Supp Fig 2A-2H). Next, we established cohorts of all genotypes treated them with tamoxifen containing food starting at 8-10 weeks of age (Fig 2A). The animals were fed the diet for 2-4 weeks and recombination of the R26-LSL-*Adar1*, reflective of removal of the stop codon and overexpression of the ADAR1 protein, was monitored by assessing peripheral blood GFP levels. This demonstrated that following tamoxifen treatment of all R26-LSL-*Adar1* had induction of GFP positive cells in the peripheral blood (Fig 2B). We deliberately tuned the tamoxifen exposure duration to generate a model where approximately half or less (on average) of the cell populations in the peripheral blood would be GFP positive based on the full length ADAR1 allele. This was done to generate a chimeric model, where cells overexpressing ADAR1 co-existed with wild-type cells in the same animal, allowing an assessment of outgrowth of ADAR1 overexpressing cells if this was an advantage conferred by gain of ADAR1 expression.

**Figure 2.**
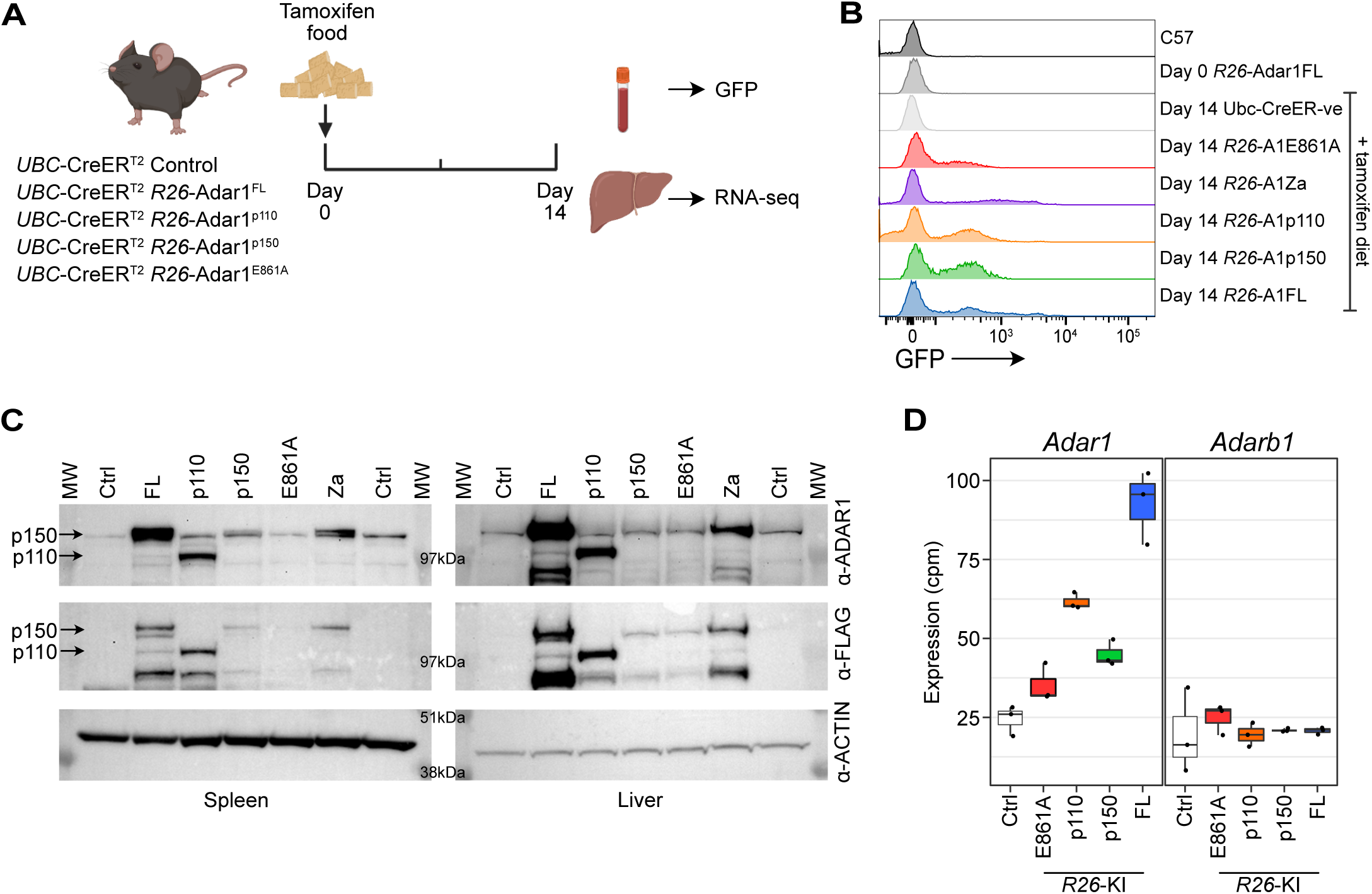
Overexpression of ADAR1 *in vivo*. **(A)** Schematic outline of *in vivo* validation experiment. **(B)** Representative flow cytometry histograms of peripheral blood leukocytes for GFP expression following 14 days *in vivo* tamoxifen treatment. **(C)** Western blot analysis of ADAR1 expression in the spleen (left panel) and liver (right panel) following 14 days *in vivo* tamoxifen treatment using anti- ADAR1 and anti-Flag antibodies. **(D)** *Adar1* and *Adarb1* expression as measured from RNA-seq of the liver following 14 days *in vivo* tamoxifen treatment (n=3-4 per genotype; note *Adar1*-Za mutation samples were not sequenced).

To determine if A-to-I editing was elevated *in vivo* we isolated samples from FL, p110, E861A and p150 animals treated with tamoxifen for 14 days. There was an induction of GFP expression, indicative of recombination and removal of the lox-Stop-lox cassette and expression of ADAR1, in the peripheral blood (Fig 2B). We confirmed *in vivo* overexpression of ADAR1 protein or the variant ADAR1 proteins using western blot analysis of both spleen (a hematopoietic organ) and liver (solid organ) at day 14 of tamoxifen administration. This demonstrated increased ADAR1 protein using both an anti-ADAR1 antibody or an ant-Flag antibody (Fig 2C). In all except the p110-mice, p150 was the dominant isoform overexpressed in the organs that were analysed. To assess the effect of overexpressing ADAR1 on the transcriptome and A-to-I editing levels, RNA was extracted from the liver at day 14 of tamoxifen administration and sequenced. We first assessed the levels of *Adar1* transcript and could demonstrate increased expression of the *Adar1* transcript in the FL, p110 and p150 samples but only a modest change in expression in the E861A samples (Fig 2D). This is consistent with the GFP and protein expression and the likely toxicity of overexpressed editing deficient ADAR1 (Fig 2B-2C). We did not see any change in the expression of the other active A-to-I editing enzyme ADAR2 because of the overexpression of ADAR1 (Fig 2D).

There was a very limited effect of ADAR1 overexpression on the cellular transcriptome, with few differentially expressed genes outside of *Adar1* itself for the FL, p110 and p150 overexpressing samples but not the E861A mutant (Fig 3A-3D; upper panel). We assessed how the overexpressed ADAR1s impacted A-to-I editing by assessing editing using a database of of 135,697 murine editing sites compiled from published databases (RADAR; (74); publications (26,29,65)) and unpublished murine datasets (JH-F, AMC and CRW). Sites required ≥50 read coverage and an editing rate of ≥0.01 (≥1%) to be considered. This demonstrated that the overexpression of full length ADAR1, p110 or p150 isoforms resulted in an increase in A-to-I editing at known sites, which appeared to correlate with the level of overexpression of the respective *Adar1* (Fig 3A-3D; lower panel). We additionally assessed the repeat site editing index (AEI; (78)) to assess the editing rates across the cohorts, allowing quantitative comparison across samples. This demonstrated that the overexpression of full length ADAR1, p110 or p150 isoforms increased the AEI (Fig 3E). The expression of the editing deficient E861A mutant does not change the AEI compared to control animals AEI (Fig 3E). The editing of the species conserved *Azin1* (p.S367>G) and *Cdk13* (p.Q103>R) were individually assessed as both are implicated in the potential oncogenic function of ADAR1 (41, 52). There was increased editing of *Azin1* at the p.S367>G site in the p150 and FL Adar1 overexpressing samples (Fig 3F), and while expression of *Cdk13* was very low in the liver tissue assessed there was evidence of increased editing in the p110 and p150 overexpressing samples (Supplemental Fig 3A). These analyses validate that the ADAR1 proteins expressed from the R26-LSL-*Adar1* alleles are functional and increase, in the case of editing proficient proteins, the levels of A-to-I editing of endogenous transcripts *in vivo*.

**Figure 3.**
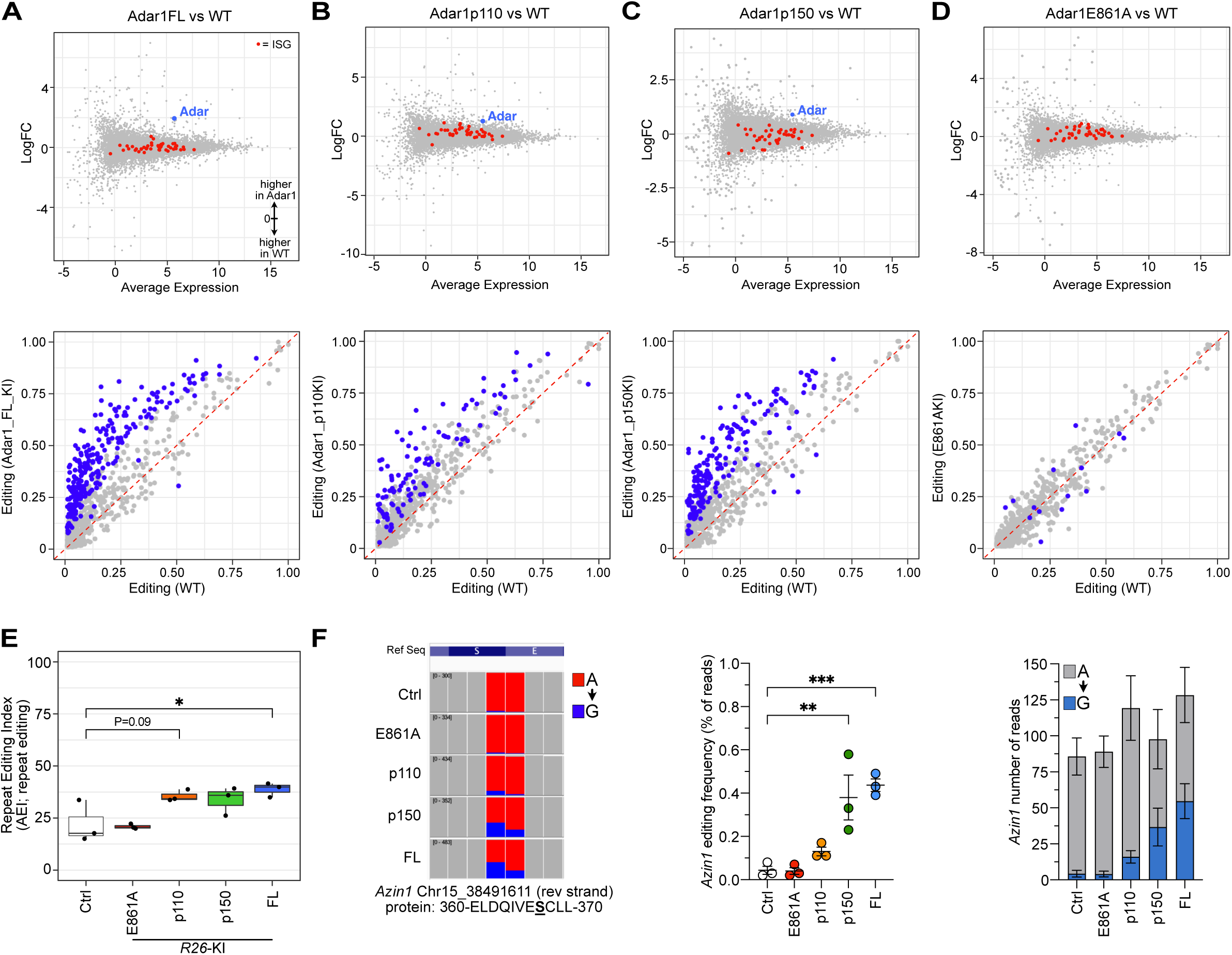
Overexpression of ADAR1 *in vivo* increased A-to-I editing levels of cellular RNA. Gene expression analysis (MA plot, upper) and A-to-I editing levels (lower) of known individual sites following 14 days *in vivo* tamoxifen treatment from **(A)** Adar1 Full length (FL); **(B)** Adar1p110; **(C)** Adar1p150; **(D)** Adar1E861A liver RNA-seq datasets compared to *Ubc*-CreER+ tamoxifen treated controls (n=3 per genotype). Red dots in upper panels represent interferon-stimulated genes. Blue dots in lower panels represent significantly different editing at individual sites between the genotypes (Jacusa statistic (likelihood ratio of two samples) >5). **(E)** The repeat editing index (AEI) from each genotype. **(F)** IGV screen shot of *Azin1* editing at the recoding p.S367G site; quantitation and statistical analysis of the editing frequency at the recoding site and average number of reads per sample for the site (expressed as mean +/- sem for each allele). \*\**P*<0.01, \*\*\**P*<0.001; Statistical comparisons using a two-way ANOVA with multiple comparisons correction using Prism software.

### Over-expression of ADAR1 *in vivo* does not modify hematopoiesis or initiate cancer

Having established that the R26-LSL-*Adar1* alleles express the expected protein products and that this resulted in an increased editing *in vivo*, we sought to determine if ADAR1 overexpression was oncogenic. We focussed our longitudinal analysis on hematopoiesis, particularly based on previous literature linking ADAR1 to both normal hematopoiesis and to hematological cancers (29,30,44,83–85). Furthermore, we could serially monitor GFP levels over time from the same cohort to monitor if ADAR1 overexpression altered normal hematopoiesis (Fig 4A).

**Figure 4.**
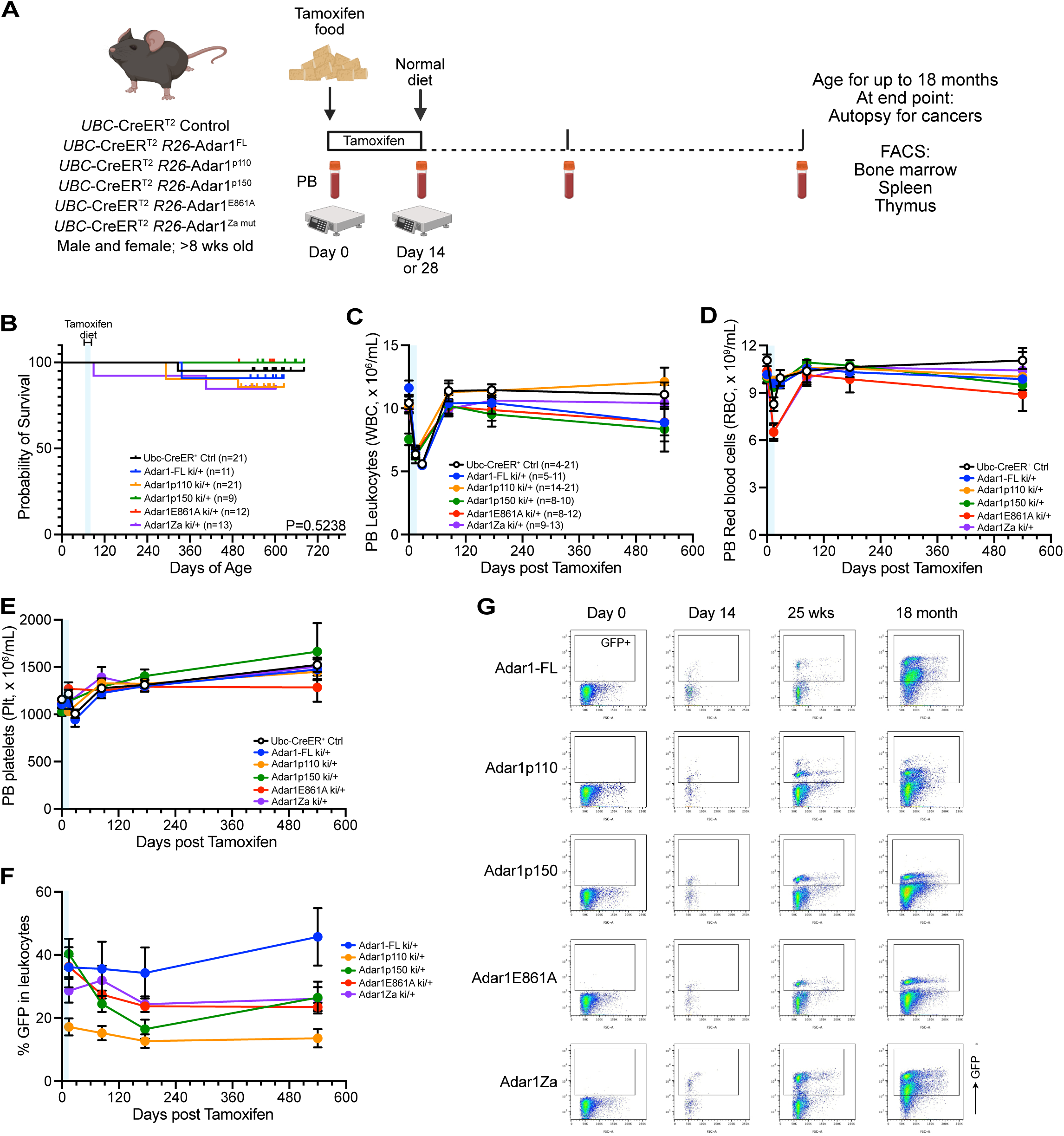
Long-term *in vivo* overexpression of ADAR1 is well tolerated. **(A)** Schematic outline of *in vivo* experiment. **(B)** Kaplan Meier survival plot of each genotype. Numbers as indicated in the inset, no significant difference in survival. Peripheral blood **(C)** leukocyte numbers, **(D)** red blood cell numbers and **(E)** platelet counts for each genotype; number per genotype indicated in panel C noting that the range indicates the minimum and maximum at any given time point per genotype. Due to restrictions during the pandemic the time points have been assigned as 0, 14/28 days, 84 days, 175 days and >580 days and the time points grouped to the closest of these for graphing. **(F)** GFP levels in the total leukocyte population. **(G)** Representative flow cytometry plots showing GFP levels in each genotype at the indicated time points. If no statistical significance indicated then no significant difference.

We established cohorts of all genotypes with littermate controls and monitored these for over 500 days (∼18 months) to determine if overexpression of ADAR1 or any of the specific isoforms impacted hematopoiesis or was sufficient to initiate cancer (Fig 4A). We did not see any impact of overexpression of ADAR1 or the different isoforms or mutant forms on long-term survival (Fig 4B). We also did not observe any elevated rates of cancer in any of the cohorts. We assessed peripheral blood (PB) parametres over time (Fig 4C-4E). When assessing overall PB indices independent of GFP levels we saw some subtle changes predominantly associated with the period where tamoxifen diet was being provided (Fig 4C-4E). After this was removed and the animals returned to a normal chow the majority of indices were comparable to the *Ubc*-CreER^T2^ R26 wild type animals (Cre+ve controls; (Fig 4C-4E). When we assessed GFP levels, as a surrogate of ADAR1 expression, we noted that the full length ADAR1 allele animals had ∼50% GFP within the granulocytes and macrophages, with slightly lower levels in the lymphoid lineage (30-40% GFP; Fig 4F-4G). The full length ADAR1 had the highest GFP expression by 18 months and then the p150 and Z*α* mutants had largely comparable GFP levels across the lineages. The E861A allele had lower intensity of GFP (Fig 2B), likely due to the selection of a level of expression of the E861A that the cells would tolerate. Unexpectedly, we also observed that the p110 total GFP levels were consistently lower across all lineages assessed (Fig 4F-4G).

At 18 months post ADAR1 overexpression we collected peripheral blood, bone marrow, spleen and thymus and undertook a detailed phenotypic analysis of a subset of the total cohort. We assessed the contribution of the ADAR1 overexpressing cells (GFP+; Fig 5) and non-ADAR1 overexpressing cells from the same animals (GFP-; Supplemental Fig 4; Supplemental Fig 8 representative FACS gating profiles). In the peripheral blood, there were no changes in the contribution of GFP+ cells to macrophages or T cells (Fig 5A). There was a reduced contribution to myeloid cells from the E861A mutant compared to the full length ADAR1 and a reciprocal increase in B cells derived from the E861A compared to the full length ADAR1 (Fig 5A). There were no changes between the full length and p110, p150 or Z*α* mutant in the myeloid or B cell populations (Fig 5A). There was no difference in the relative contributions of GFP- cells to all lineages assessed in the peripheral blood (Supplemental Fig 4A). Total bone marrow cellularity, independent of GFP, was comparable across all cohorts (Fig 5B). In the bone marrow there was no changes in the contribution of GFP+ cells to the B or T lymphoid cells (Fig 5C). The ADAR1p150 overexpressing cells had a lower contribution to the neutrophil/granulocyte lineage in the bone marrow compared to the full length, p110 and E861A alleles that was not apparent in the peripheral blood. Note we did not include erythroid cells in this analysis as the red blood cells enucleate and lose GFP signal, making definitive assignment to GFP+/GFP- populations confounding. Peripheral blood erythroid indices were comparable across all genotypes (Fig 4D). We assessed contribution to the hematopoietic stem and progenitor populations (Fig 5D-5E). There was no change in contributions to the phenotypic long-term (LT-HSC) or short-term hematopoietic stem cell (ST-HSC) population within the lineage^-^cKit^+^Sca-1^+^ (LKS+) population (Fig 5F). There was an increased proportion of the progenitor enriched LKS^-^ in the E861A and Z*α* mutants compared to the *Adar1* full length allele, however, all of the specific populations within the LKS- population had a comparable contribution from the GFP+ cells (Fig 5G). The spleen (Fig 5H) and thymus (Fig 5I) were largely comparable across the different genotypes. Collectively these analyses indicate that the long-term overexpression of ADAR1, p110, p150, E861A or Z*α* is well tolerated and results in only modest effects on hematopoiesis.

**Figure 5.**
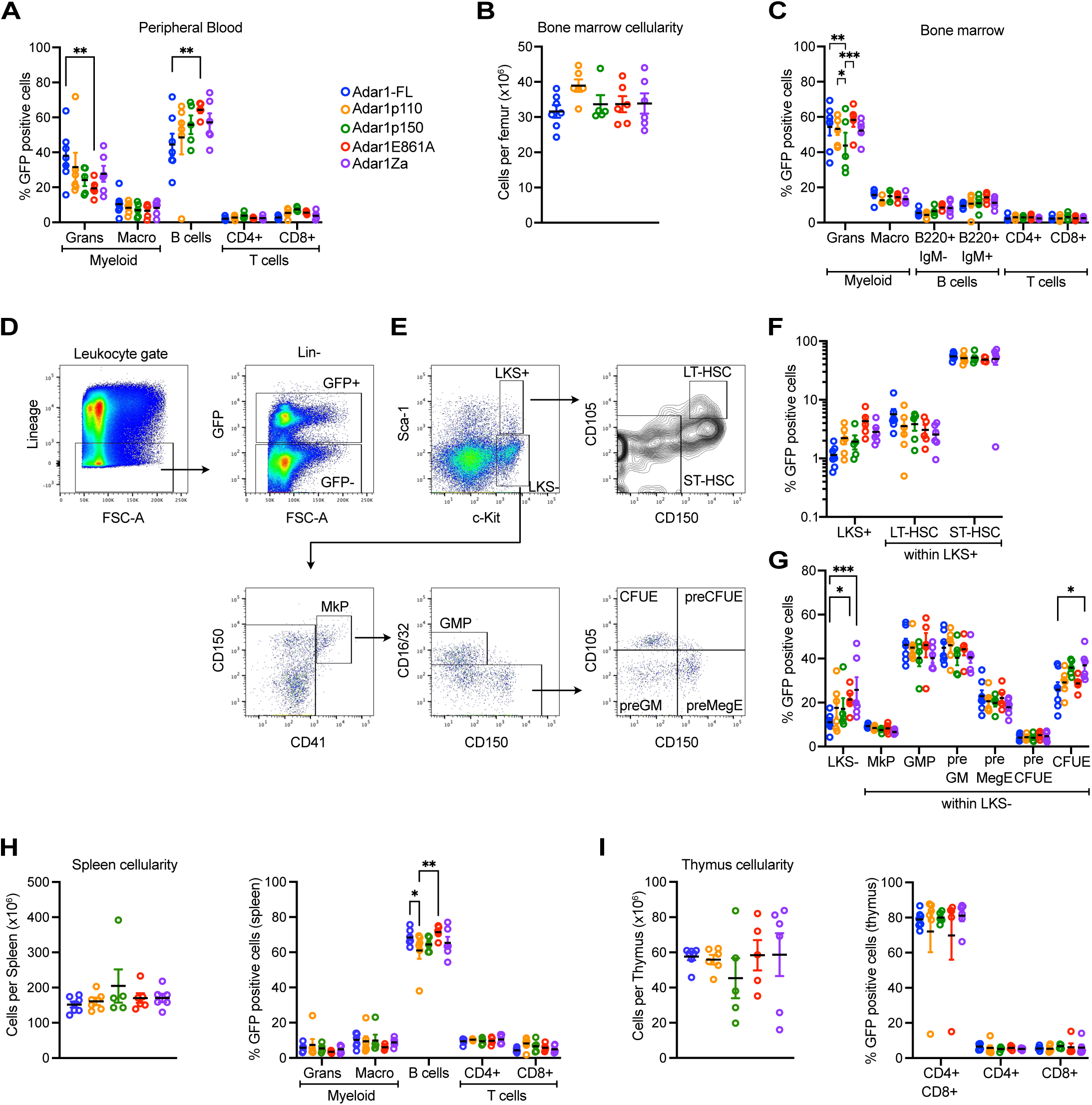
Modest changes in hematopoiesis with overexpression of ADAR1. **(A)** The percentage contribution of peripheral blood GFP+ cells to each indicated lineage across each genotype. **(B)** Total bone marrow cellularity per femur. **(C)** The percentage contribution of GFP+ cells in the bone marrow to each indicated population. **(D-E)** representative flow cytometry plots used to assess the hematopoietic stem and progenitor compartment. **(F)** The percentage contribution of GFP+ cells in the bone marrow to the lineage-cKit+Sca1+ (LKS+) population and the long-term and short-term hematopoietic stem cell populations (contained within the LKS+ fraction). **(G)** The percentage contribution of GFP+ cells in the bone marrow to the lineage-cKit+Sca1- (LKS-) population and the megakaryocyte progenitors (MkP), granulocyte macrophage progenitors (GMP), pre-GM, pre- Megakaryocyte erythroid progenitors (preMegE), pre colony forming unit erythroid (preCFU-E) and CFU-E populations (contained within the LKS- fraction). **(H)** Total cellularity of the spleen and contribution of the GFP+ cells to the indicated cell populations. **(I)** Total thymus cellularity and contribution of the GFP+ cells to the indicated cell populations. Each circle indicates an individual animal; \**P*<0.05, \*\**P*<0.01, \*\*\**P*<0.001; Statistical comparisons using a two-way ANOVA with multiple comparisons correction using Prism software.

Coupled with the longitudinal analysis described above we completed autopsies on all animals to assess for any changes in non-hematological organs and for detection of any potential cancers. A total of 4 mice across all cohorts required euthanasia prior to the 18-month time point. Two of the four mice were determined to have developed cancer: colon cancer in one *Ubc*-CreER^T2Tg/+^ R26-*Adar1^ki/+^* model and lymphoma in one *Ubc*-CreER^T2Tg/+^ R26-*Adar1p150^ki/ki^* model. The remaining two mice (*Ubc*-CreER^T2Tg/+^ R26-wildtype and *Ubc*-CreER^T2Tg/+^ R26-*Adar1^ki/+^*) had inconclusive autopsies, with no tumors apparent so the cause of death was not determined to be related to cancer. Kaplan-Meier survival analysis demonstrated no significant difference in the survival between the control and ADAR1 isoform overexpressing mouse models (Fig 3B). A p110 overexpressing animal had very elevated myeloid cell contribution to the peripheral blood at 18 months of age, but upon further analysis this was being contributed to by both the GFP+ (ADAR1 overexpressing) and GFP- cells suggesting that this was not a phenotype that directly resulted from ADAR1p110 overexpression (Supplemental Fig 5). Collectively, these results demonstrate that the long-term overexpression of ADAR1, or its individual isoforms, *in vivo* is well tolerated and not sufficient to initiate tumor formation as a single lesion. This indicates that overexpression of ADAR1 in isolation was not sufficient to act as an initiator of tumorigenesis.

### Endogenous ADAR1 and A-to-I editing increase upon transformation

We further sought to explore the role of ADAR1 overexpression in solid tumor formation. We first tested if endogenous *Adar1* and A-to-I editing increased upon cell transformation in murine cells, as demonstrated in human tumors (40–44). We used a murine osteoblast immortalisation protocol that takes primary long bone derived osteoblasts and upon engineered deletion of p53 (*R26*-CreER^T2^ *Tp53^fl/fl^* derived cells) these cells immortalise and have been demonstrated to generate osteosarcoma when transplanted *in vivo* (63,86–88). We collected cells at day 7, 14 and 21 post tamoxifen addition and completed RNA-seq to assess *Adar1* levels and quantitate A-to-I editing levels (Fig 6A). We see a progressive increase in endogenous *Adar1* levels from day 7 to 21, with an ∼3 fold increase by day 21 when the cells have become p53 deficient (Fig 6B; Supplemental Fig 6A). We do not see an increase in the expression of *Adarb1*, encoding ADAR2, the other enzymatically active ADAR in mammals (Fig 6B). The increase in *Adar1* was accompanied by an increase in A-to-I editing, whether assessed at individual recoding sites in *Azin1*, across known editing sites in mouse samples or using the AEI index, demonstrating that the increased transcript was resulting in an increased level of A-to-I editing (Fig 6D-6F)(78). *Cdk13* had a high basal editing level in this cell type so we could not discern significant changes. Interestingly, accompanying the transition of the cells to being p53 deficient and increased *Adar1* expression was an increased expression of the transcriptional program associated with Type I interferon (Fig 6C, Supplemental Fig 6B). This is a very similar signature associated with a loss of ADAR1 or a loss of ADAR1 mediated A-to-I editing but has also been reported to arise in *p53^-/-^* MEFs due to sensing of endogenous mitochondrial dsRNA (29,30,89). There is a co-ordinated increase in the expression of these transcripts independent of infection, indicating that immortalisation/transformation induced by a loss of p53 is accompanied by an activation of the Type I interferon transcriptome, which includes *Adar1*.

**Figure 6.**
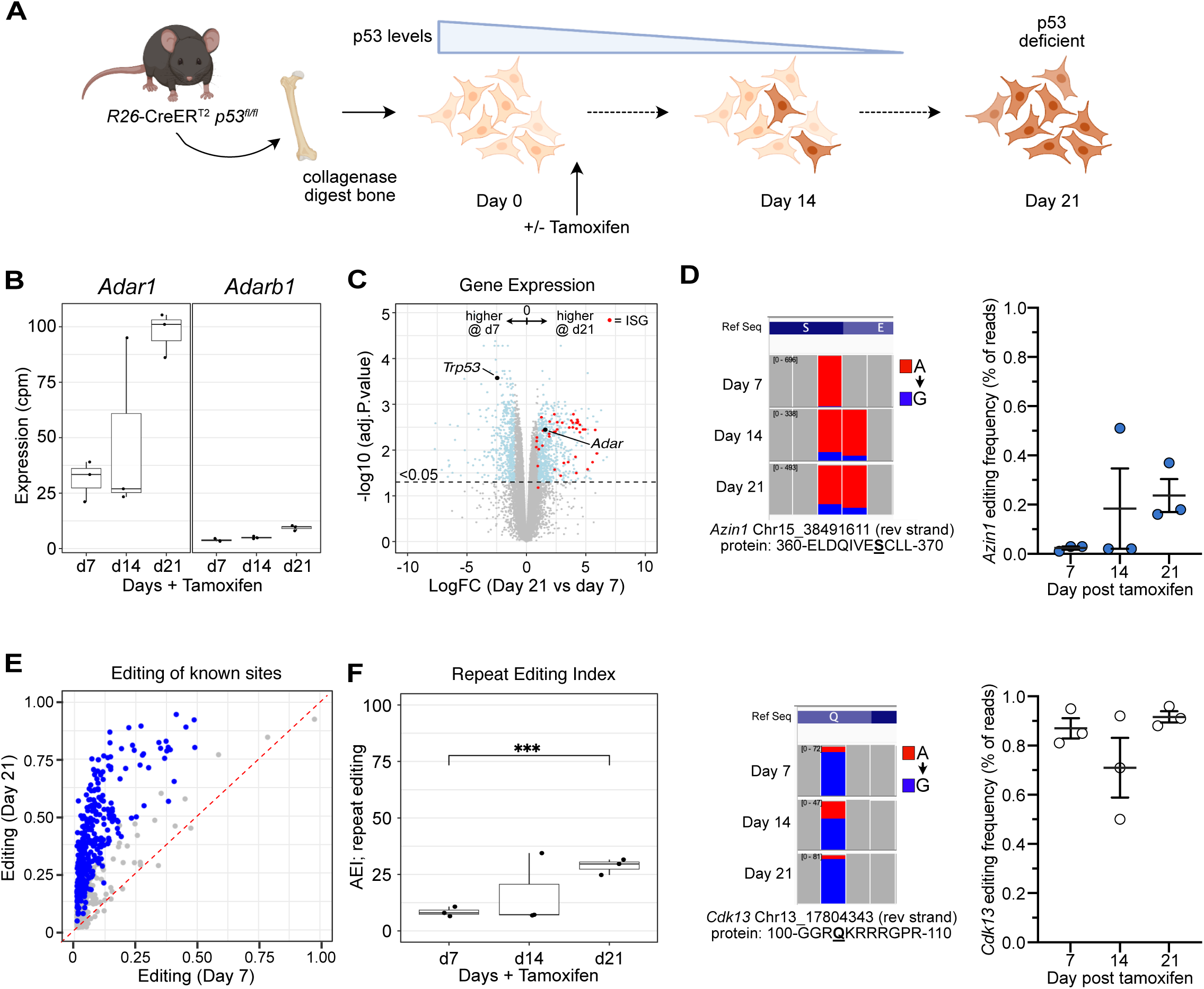
*Adar1* expression and editing activity increases during cellular immortalization. **(A)** Schematic outline of *in vitro* experiment. Long bone osteoblasts were isolated from *R26*-CreER^T2^ *Trp53^fl/fl^* mice and cultured with tamoxifen to induce deletion of p53. Cells were collected at day 7, 14 and 21 for analysis. Osteoblasts were isolated from three animals and cultured separately (biological replicates). **(B)** Expression (counts per million) of *Adar1* and *Adarb1* at day 7, 14 and 21 as determined by RNA-seq at each time point. **(C)** MA plot of gene expression comparing day 7 (p53 still expressed) and day 21 (p53 deficient) with significantly different genes indicated in blue and interferon stimulated genes (ISGs) indicated in red. **(D)** IGV screen shot of *Azin1* (upper) and *Cdk13* (lower) editing at the indicated recoding sites; quantitation of the editing frequency at each site (expressed as mean +/- sem for each allele). The Alu/repeat editing index (AEI) from each timepoint derived from the RNA-seq. **(E)** A-to-I editing levels of known individual sites comparing editing levels at day 21 (y axis) to day 7 (x axis) following tamoxifen treatment. Blue dots represent significantly different editing at individual sites between the genotypes (Jacusa statistic (likelihood ratio of two samples) >5). **(F)** The repeat editing index (AEI) at each time point calculated from the RNA-seq dataset. \*\*\**P*<0.001; Statistical comparisons using a two-way ANOVA with multiple comparisons correction.

### ADAR1 overexpression does not accelerate cancer formation initiated by loss of tumor suppressor genes

Having established that overexpression of ADAR1, or either the p110 or p150 isoforms individually, are not sufficient to induce tumor formation as sole lesions *in vivo* we sought to determine if ADAR1 overexpression would cooperate to accelerate tumour formation dependent upon other mutations. To this end we made use of a highly characterised autochthonous model of osteosarcoma, the most common cancer of bone (55, 62). This model is based on the osteoblast progenitor restricted deletion of *Tp53* and retinoblastoma protein (*Rb*) (55, 90). This model yields a fully penetrant osteosarcoma model, with a subset of animals also developing metastasis, that replicates to the cardinal features of human osteosarcoma (55, 91). Additionally, it is known that there is a physiologically essential function for ADAR1 in osteoblasts *in vivo*, based on the phenotypes associated with the loss of function analysis in mice (92) demonstrating that ADAR1 has a function in this cell type normally. Recent analysis of paediatric cancers demonstrates that A-to-I editing can be observed in human osteosarcoma (93).

We crossed the *Osx1*-Cre *p53^fl/fl^ pRb^fl/fl^* to the R26-LSL-*Adar1* alleles to generate cohorts of all possible genotypes, except those with the Z*α* mutation which were not used in the solid tumor studies (Fig 7A). It should be noted that the *Osx1*-Cre transgene used for these models is active in all pre-osteoblast populations, including during development, and that the Cre activity was not being regulated temporally in this model (55, 62). We used either *Osx1*-Cre *p53^fl/fl^ pRb^fl/+^* or *Osx1*-Cre *p53^fl/fl^ pRb^fl/fl^* genotypes as these have comparable latency and disease manifestations (55, 62). We set aside cohorts of each genotype and allowed them to age to assess if overexpression of ADAR1 altered the tumour latency, number of primary tumours arising, metastatic potential or tumor subtype. We compared these to littermate *Osx1*-Cre *p53^fl/fl^ pRb^fl/+^* or *Osx1*-Cre *p53^fl/fl^ pRb^fl/fl^* genotypes that were aged in parallel and historical datasets from this model in the same facility. Mice were monitored for tumor formation, as determined by animal facility staff. The animal facility staff were unaware of the *Rosa26* genotype so were not likely to introduce selection bias into the detection of tumors. Following detection of a tumour mass, the animals were closely monitored and then euthanased when they reached ethical endpoint. This was followed by autopsy, tumor resection and isolation, and pathology was performed to characterise the tumor phenotype.

**Figure 7.**
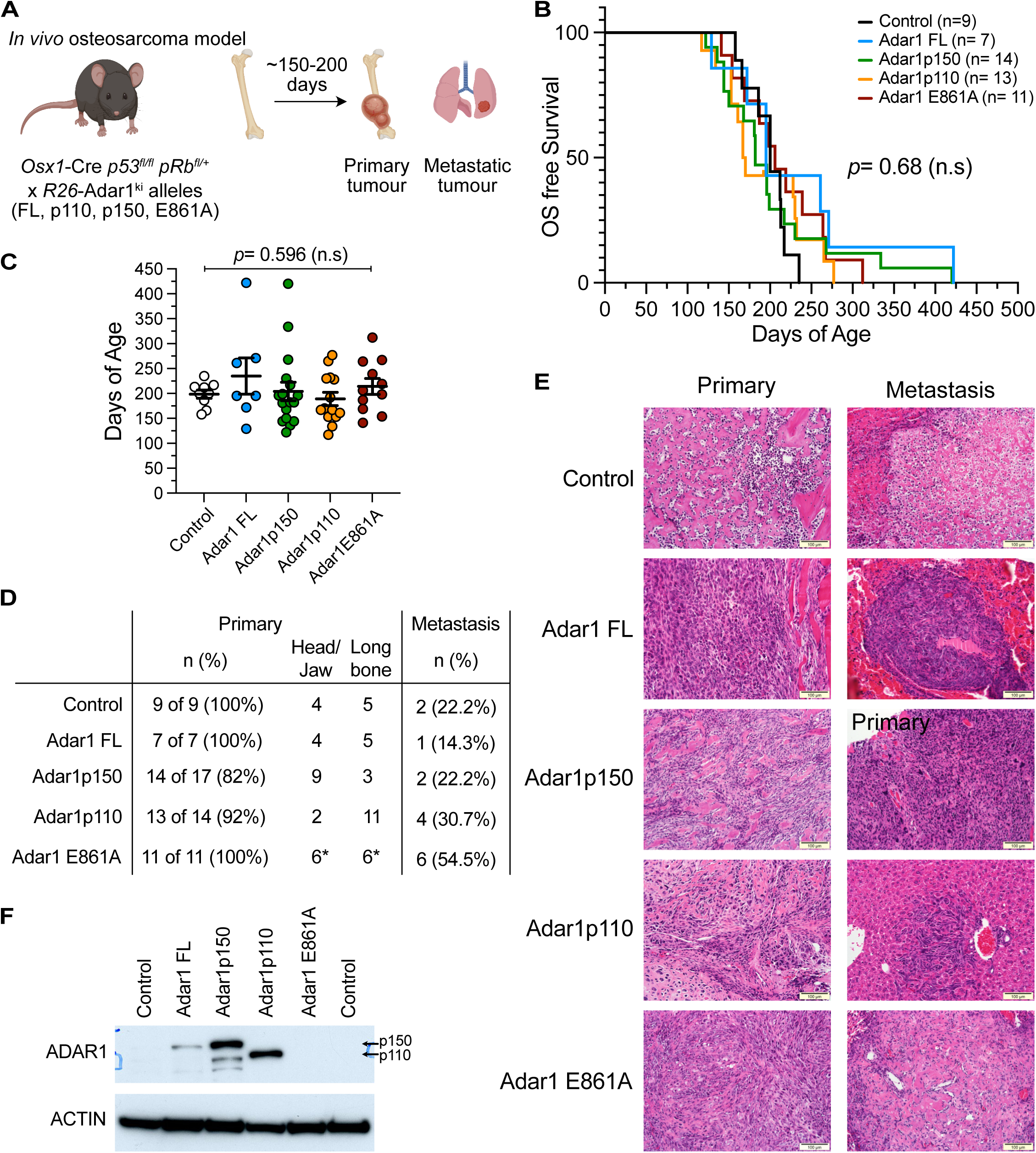
Overexpression of ADAR1 does not accelerate or modify osteosarcoma behavior *in vivo*. **(A)** Schematic outline of *in vivo* osteosarcoma model. **(B)** Kaplan Meier survival plot of each genotype. Numbers as indicated in the inset, no significant difference in survival. Number of animals per genotype indicated in inset. **(C)** Median days of age of each genotype, same cohort as represented in the KM plot; assessed by two-way ANOVA with multiple comparison correction. **(D)** Analysis of primary tumor location and metastatic spread in each genotype. **(E)** Representative histology of primary and metastatic lesions from each indicated genotype. The osteosarcoma model generates a fibroblastic osteosarcoma. **(F)** Western blot of ADAR1 (anti-ADAR1 antibody) in whole tumor pieces derived from the indicated genotypes.

Comparative analysis between the control and ADAR1 overexpressing cohorts determined no significant difference in overall survival (Fig 7B-7C; Supplemental Fig 7A-7B). All animals developed osteosarcoma with a median survival of 172.5 days (*Osx1*-Cre *p53^fl/fl^ pRb^fl/+^* ^or *fl/fl*^ R26-*Adar1p150^ki/+^*) through to 228 days (*Osx1*-Cre *p53^fl/fl^ pRb^fl/+ or fl/fl^* R26-*Adar1p110^ki/+^*) (Fig 7C). Median survival for control *Osx1*-Cre *p53^fl/fl^ pRb^fl/+^* was 212 days, comparable to the historical data from large cohorts in the same facility demonstrating a median survival between 127-206 days in *Osx1*-Cre *p53^fl/fl^ pRb^fl/+^* and *Osx1*-Cre *p53^fl/fl^ pRb^fl/fl^* mice (62, 63). The primary tumors were phenotypically similar across all cohorts, with smooth rounded calcified masses originating from bone. Metastasis were observed primarily within lung and liver tissue, and there was evidence of spread within lymphatic tissue and consistent with both human osteosarcoma and the dissemination patterns reported in this model (Fig 7D) (55,62,63). Histology was performed on sections of primary and metastatic tumors from each cohort (Fig 7E). This confirmed that the tumors were primarily fibroblastic osteosarcoma, as expected with this model (55, 62). We then compared the rate of metastasis in the cohorts to determine if elevated ADAR1 expression resulted in a change in metastatic frequency. A higher metastatic rate would imply more aggressive and rapidly spreading tumours and indicate an advantageous role for gain of ADAR1 expression in tumor progression. Z-score analysis of the metastatic frequency across the cohorts found no significant increase in the rate of metastasis following ADAR1 overexpression compared to the control cohort. We checked for expression of ADAR1 in the tumors by Western blot and could show higher expression of FL, p150 and p110 in the overexpressing models relative to control tumors where ADAR1 was below the limits of detection. Consistent with previous data suggesting overexpressing an editing-dead protein is not well tolerated, we could not detect increased ADAR1 in the E861A mice (Fig 7F). Collectively this data indicates that while ADAR1 overexpression has been reported in many human cancer types, we do not find experimental evidence to directly support a role for elevated ADAR1 expression or activity in either the initiation or progression of cancer *in vivo* in the mouse.

## DISCUSSION

The large-scale analysis of the transcriptome of diverse cancers has led to the identification of elevated A-to-I RNA editing, and the writer of this pervasive epitranscriptomic mark ADAR1, as generalizable features of human cancer (7,21,40). In parallel, results from functional genetic screening and gene specific analysis have identified that loss of ADAR1 as a potent sensitizer to immune checkpoint blockade therapy (8,10,12). These data have led to intensive efforts to develop inhibitors of ADAR1 for oncology. Whilst these findings of increased ADAR1 levels and activity in cancer and the genetic sensitivity of subsets of cancer to loss of ADAR1 have been broadly supported, the role of elevated ADAR1 in tumor initiation and progression have only been tested in limited settings. A more refined understanding of the role of ADAR1 in tumor initiation and maintenance will be important to understand at which stage in the formation of a cancer ADAR1 expression is elevated and for insight into the contribution of A-to-I editing to tumor evolution. Here we have sought to directly test the effects of elevated ADAR1 levels *in vivo*, using well characterised mouse models, to understand the role that ADAR1 plays in oncogenesis.

We established a series of knock-in mouse models that would allow controlled expression of ADAR1 (both p110 and p150), the individual p110 or p150 isoforms *in vivo* in response to Cre mediated recombination. We also established two additional lines, an E861A mutant that yields an editing dead p110/p150 and acts as a control (29), and a N175A/Y179A mutant in the p150 specific Z*α* domain (53, 79). When we initiated our studies there was relatively scant information as to the function of the Z*α* domain, however a number of recent studies have significantly increased understanding of this p150 unique domain (79, 81). We utilized a broadly active, somatically inducible Cre (*Ubc*-CreER) to mediate widespread expression of ADAR1 across the different tissues and organs of the mouse, and both male and female mice were used for all analysis. Short- and long-term analysis and monitoring did not demonstrate an increased frequency of cancers in these mice.

We deliberately chose a strategy to yield a chimeric animal where approximately half or less of the cells in the peripheral blood (based on the co-expressed GFP marker) would be overexpressing ADAR1 and the remaining cells had physiological ADAR1 levels (GFP-ve, not overexpressing). We reasoned that if ADAR1 was advantageous, even if not to the point of being sufficient to initiate cancer as a sole driver, we may see an advantage reflected with the progressive outgrowth of ADAR1/GFP+ overexpressing cells with time. We achieved an approximately 3-fold increase in the expression of full length *Adar1* transcript from the *Rosa26* alleles (Fig 2D), comparable to the level of increased *ADAR* transcript reported across a range of human cancers compared to controls (43) and of the endogenous murine *Adar1* following immortalization by loss of p53 (Fig 6B). Over a period of 18 months monitoring, we did not observe increased levels of GFP in any of the different cohorts. This result suggests that the ADAR1/GFP overexpressing cells were not conferred with a significant advantage from this modification, at least within the hematopoietic system. Whilst the GFP monitoring was restricted to the cells and organs of the hematopoietic system (peripheral blood, bone marrow, spleen, thymus), upon autopsy we assessed all animals for any evidence of changes in organs potentially indicative of cancer. We did not find any macroscopic evidence of reproducible changes in any organs that would be consistent with ADAR1 overexpression, or its individual isoforms p110 or p150, leading to formation of tumors *in vivo*. It has been demonstrated using analogous models that the C-to-U base editor APOBEC3 will induce tumors in mice following overexpression *in vivo*, demonstrating that such systems will yield *in vivo* cancer if the overexpressed protein can function as an oncogene (94–96). Based on the two models we proposed, we did not find evidence to support the conclusion that ADAR1 was functioning as an oncogene.

In parallel, we sought to determine if the overexpression of ADAR1 would co-operate to promote tumor formation or progression and metastasis in a tumor initiated by separate genetic event(s). We can recapitulate an increase in *Adar1* expression and A-to-I editing levels in a reductionist model of cellular immortalisation, in this instance the immortalisation of primary osteoblasts following loss of p53. This demonstrates that in mouse cells transforming there is a similar change in ADAR1 and A-to-I editing as reported across a range of human tumors (40, 42). Perhaps most interestingly, this model system demonstrated that as cells transitioned to a p53 deficient state they also had an elevated interferon stimulated gene (ISG) transcriptional signature. This was unexpected as loss of ADAR1, or specifically its editing activity, also activates a similar transcriptional response (29, 30). In this setting, the elevated *Adar1* transcript and A-to-I editing are most likely secondary to the activation of the ISG signature, given *Adar1p150* is a well characterised ISG and there is increasing evidence that *Adar1p110* is also induced to some extent by Type I interferon (35,97–99). We found a 2-3 fold elevation in the expression of endogenous *Adar1* as the osteoblastic cells lost p53 and immortalised, consistent with the range in increased *ADAR* reported in human cancers (43). These data are most consistent with that reported in human breast cancer, where elevated *ADAR1* was associated with the amplification of 1q and inflammation (42). Our results align with those reported following analysis of the immunogenicity of *p53^-/-^* MEF derived cellular dsRNA, which demonstrated that this genotype generated immunogenic dsRNA of mitochondrial origin (89). These data support the model where elevated ADAR1 in cancers is a secondary response to tumor formation and the changes in the environment, both the physical environment and the transcriptional landscape of the tumor.

We then made use of a well characterised osteosarcoma model that develops at a highly reproducible latency and can undergo spontaneous metastatic spread to test the effects of ADAR1 overexpression in co-operation with loss of tumor suppressor genes (55). This allowed an assessment of the effect of overexpression of ADAR1 on both primary and metastatic tumor behaviour *in vivo*. The alternative model we hypothesised, ADAR1 as a passenger (Fig 1A), would predict that the overexpression of ADAR1 may be advantageous to the tumor and provide a means to suppress immune responses to cellular dsRNA. A prediction of this model is that the overexpression of ADAR1 could either shorten the latency of primary tumor formation or facilitate a greater frequency/extent of metastatic spread. We did not observe either effect in the osteosarcoma model, a tumor initiated by the loss of *Trp53*. In these experiments we did not see evidence for modulation of the latency or metastatic potential within the cohort size assessed. We have only completed a macroscopic assessment and this is a limitation of the study, as we would not appreciate any changes in tumour infiltrating cells that the overexpression for ADAR1 may modulate. Whilst this is an important caveat of our work, in the context of the model we have used which is immune competent, the effects of overexpression of ADAR1 are not sufficient to modulate the survival time meaningfully.

Collectively these studies demonstrate that overexpression of ADAR1 full length, or its individual isoforms ADAR1p110 and ADAR1p150, is not sufficient to initiate cancer *in vivo* in the mouse. We find long-term tolerance to an elevated level of ADAR1, p110 or p150 and that this is overall well tolerated *in vivo*. We can demonstrate that Adar1 levels increase as mouse cells undergo immortalisation induced by the loss of a tumor suppressor gene and that this is associated both with an increase in A-to-I editing and a more general activation of interferon stimulated gene expression. It is known that A-to-I editing is increased in the context of interferon treatment, which can also induce Z-RNA (8, 53). In the context of an osteosarcoma model that mirrors human cancer, overexpression of ADAR1 did not significantly modify any of the parameters measured. We conclude that ADAR1 overexpression and the increased A-to-I editing reported in many human cancer types is most likely a consequence of tumor formation and that it is not sufficient to initiate cancer as a single event.

## DATA AVAILABILITY

RNA-seq datasets generated for this study are available in NCBI GEO (Accession number: GSE221628)

## SUPPLEMENTARY DATA

### Figures

**Supplemental Figure 1.**
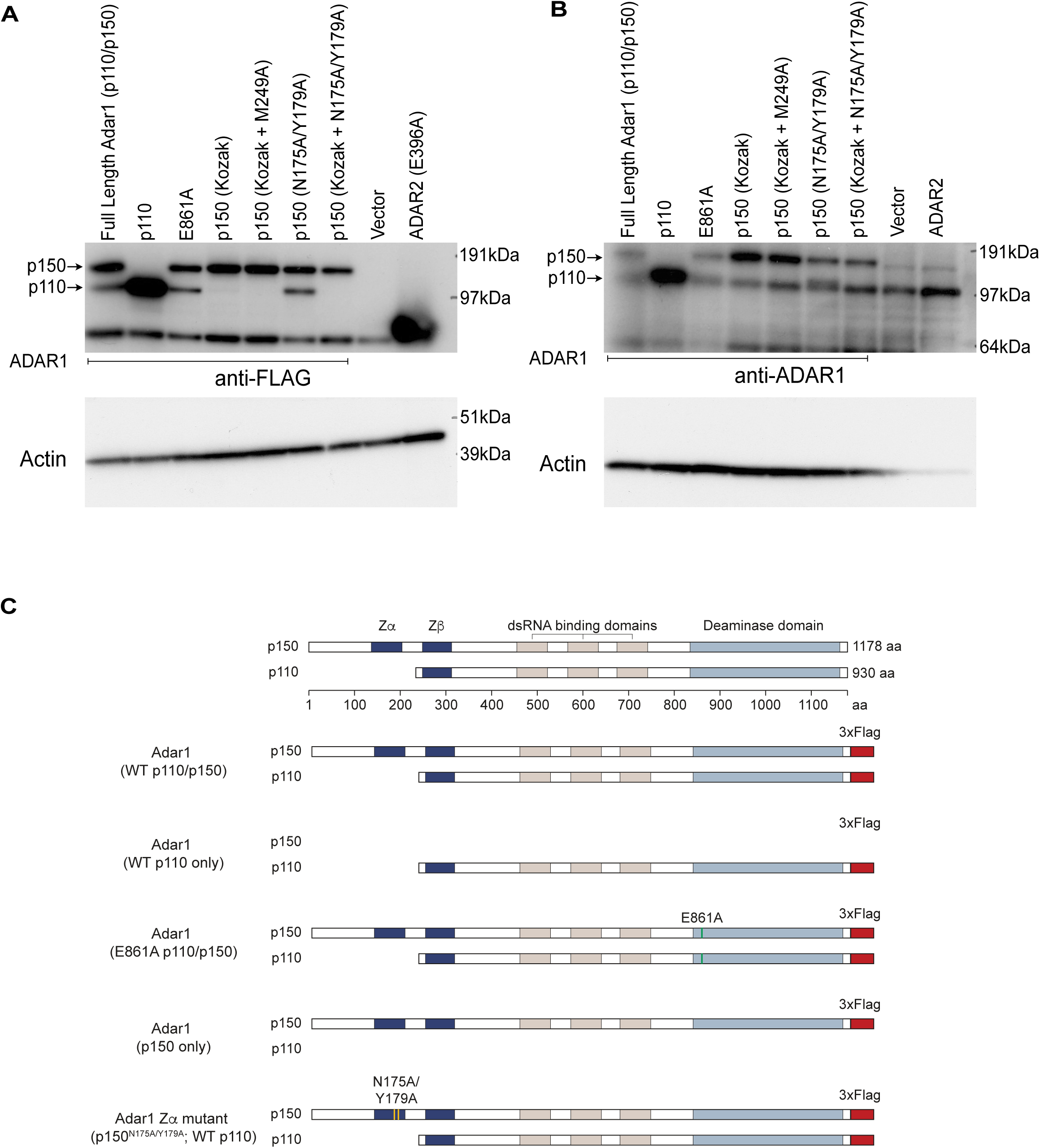
Validation of Adar1 cDNAs. Expression of the indicated murine *Adar1* cDNA using lentivirus in the murine stromal cell line Kusa4b10 and western blot using **(A)** anti-Flag or **(B)** anti-ADAR1 antibody. Text in brackets indicates any modification to the *Adar1* cDNA. *Adar2* included as a control. **(C)** Graphical representation of the different protein products expected from each allele.

**Supplementary Figure 2.**
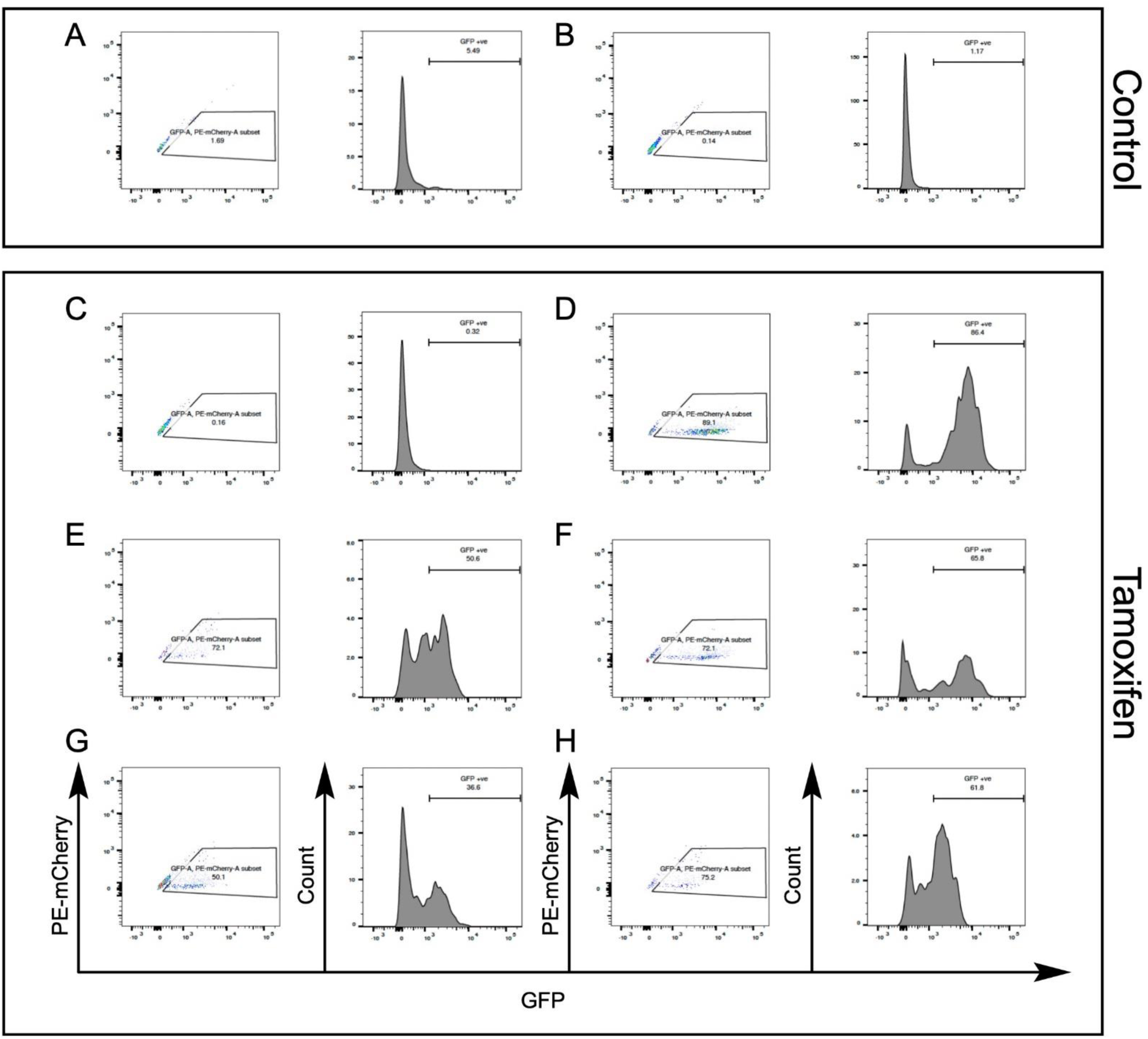
Expression of GFP in adult tail fibroblasts derived from *Ubc*-CreER mice of the indicated genotype. Flow cytometry analysis of adult tail fibroblasts: Untreated **(A)** *Ubc*- CreER-ve *R26*-*Adar1^ki/ki^* and (B) *Ubc*-CreER^tg/+^ *R26*-*Adar1^ki/ki^* cell lines. Panel **(C-H)** are tail fibroblasts treated with 400nM 4-hydroxy tamoxifen for 7 days: **(C)** *Ubc*-CreER-ve *R26*-*Adar1^ki/ki^* (Cre negative; from panel A); **(D)** *Ubc*-CreER^tg/+^ *R26*-*Adar1^ki/ki^* (Cre+ve from panel B); **(E)** *Ubc*-CreER^tg/+^ *R26*- *Adar1p150^ki/ki^*; **(F)** *Ubc*-CreER^tg/+^ *R26*-*Adar1-Za^ki/ki^*; **(G)** *Ubc*-CreER^tg/+^ *R26*-*Adar1p110^ki/ki^* and **(H)** *Ubc*- CreER^tg/+^ *R26*-*Adar1E861A^ki/ki^*.

**Supplementary Figure 3.**
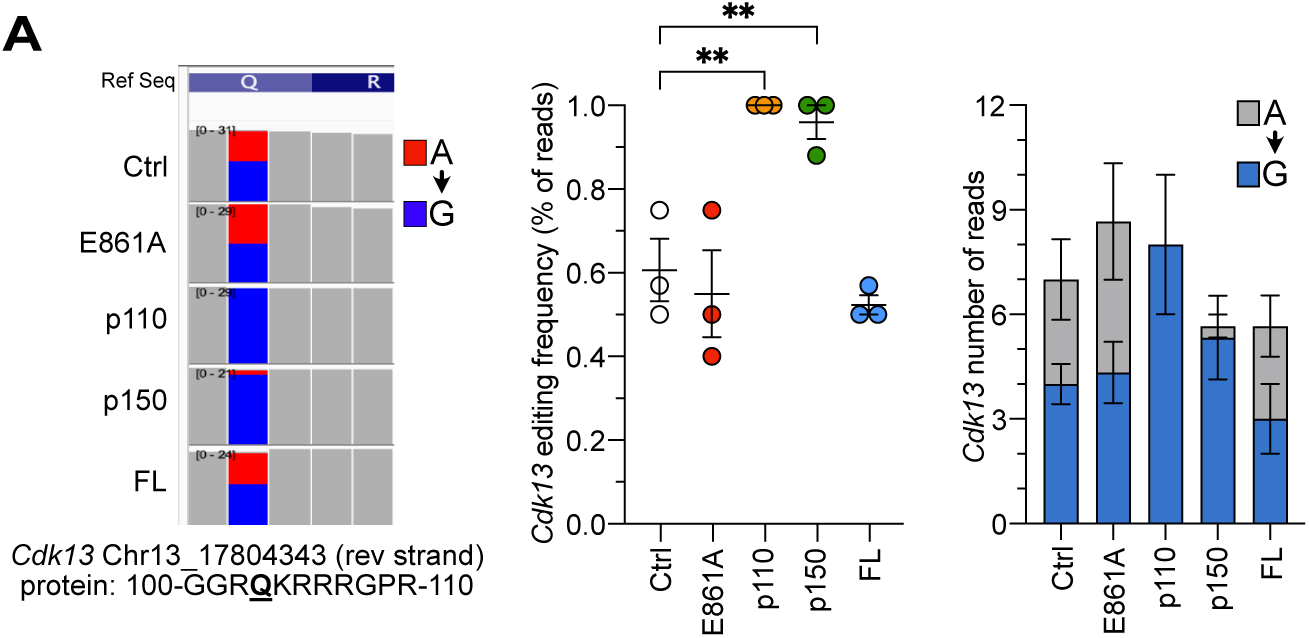
Editing of *Cdk13* recoding site in liver samples following 14 days of tamoxifen treatment. **(A)** IGV screen shot of *Cdk13* editing at the recoding p. Q103>R site; quantitation and statistical analysis of the editing frequency at the recoding site and average number of reads per sample for the site (expressed as mean +/- sem for each allele). \*\**P*<0.01, \*\*\**P*<0.001; Statistical comparisons using a two-way ANOVA with multiple comparisons correction using Prism software.

**Supplementary Figure 4.**
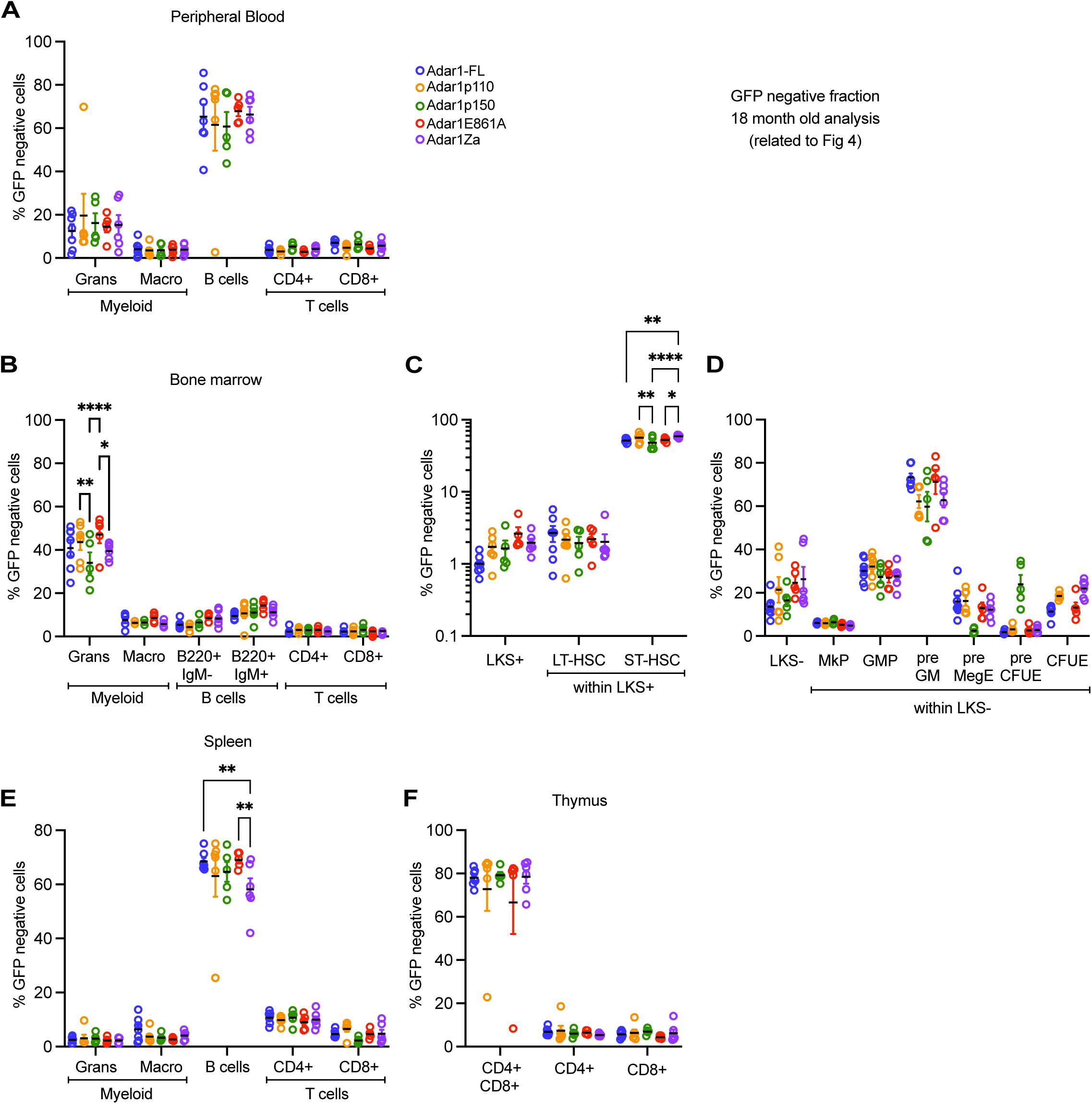
Analysis of the contribution of the GFP-ve cells from the analysis of 18 month old animals. **(A)** The percentage contribution of peripheral blood GFP- cells (non-ADAR1 overexpressing cells) to each indicated lineage. **(B)** The percentage contribution of GFP- cells in the bone marrow to each indicated population. **(C)** The percentage contribution of GFP- cells in the bone marrow to the lineage-cKit+Sca1+ (LKS+) population and the long-term and short-term hematopoietic stem cell populations (contained within the LKS+ fraction). **(D)** The percentage contribution of GFP- cells in the bone marrow to the lineage-cKit+Sca1- (LKS-) population and the megakaryocyte progenitors (MkP), granulocyte macrophage progenitors (GMP), pre-GM, pre-Megakaryocyte erythroid progenitors (preMegE), pre colony forming unit erythroid (preCFU-E) and CFU-E populations (contained within the LKS- fraction). **(E)** Contribution of the GFP- cells to the indicated cell populations. **(F)** Contribution of the GFP- cells to the indicated cell populations. Each circle indicates an individual animal; \**P*<0.05, \*\**P*<0.01, \*\*\**P*<0.001; \*\*\*\**P*<0.0001; Statistical comparisons using a two-way ANOVA with multiple comparisons correction using Prism software.

**Supplementary Figure 5.**
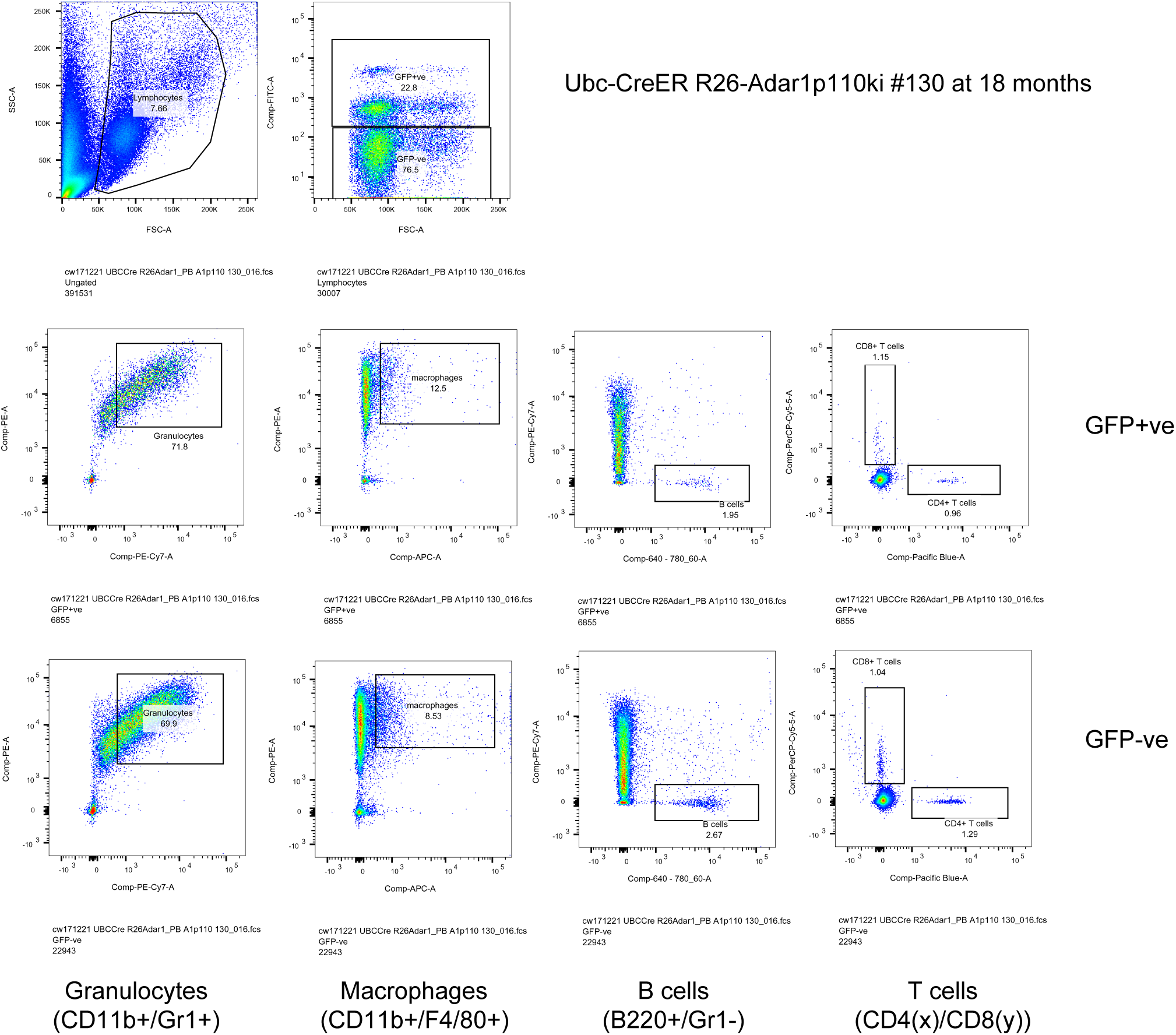
FACS analysis of peripheral blood from *Ubc*-CreER R26-Adar1p110 mouse #130 at 18 months post tamoxifen. Analysis of haemolysed peripheral blood leukocytes in *Ubc*-CreER R26-Adar1p110 mouse #130 at 18 months post tamoxifen. Top panel shows gating of leukocytes (FSC v SSC) and the GFP gating from the total leukocyte population. The lineage contribution from the GFP+ve and GFP-ve are then shown.

**Supplementary Figure 6.**
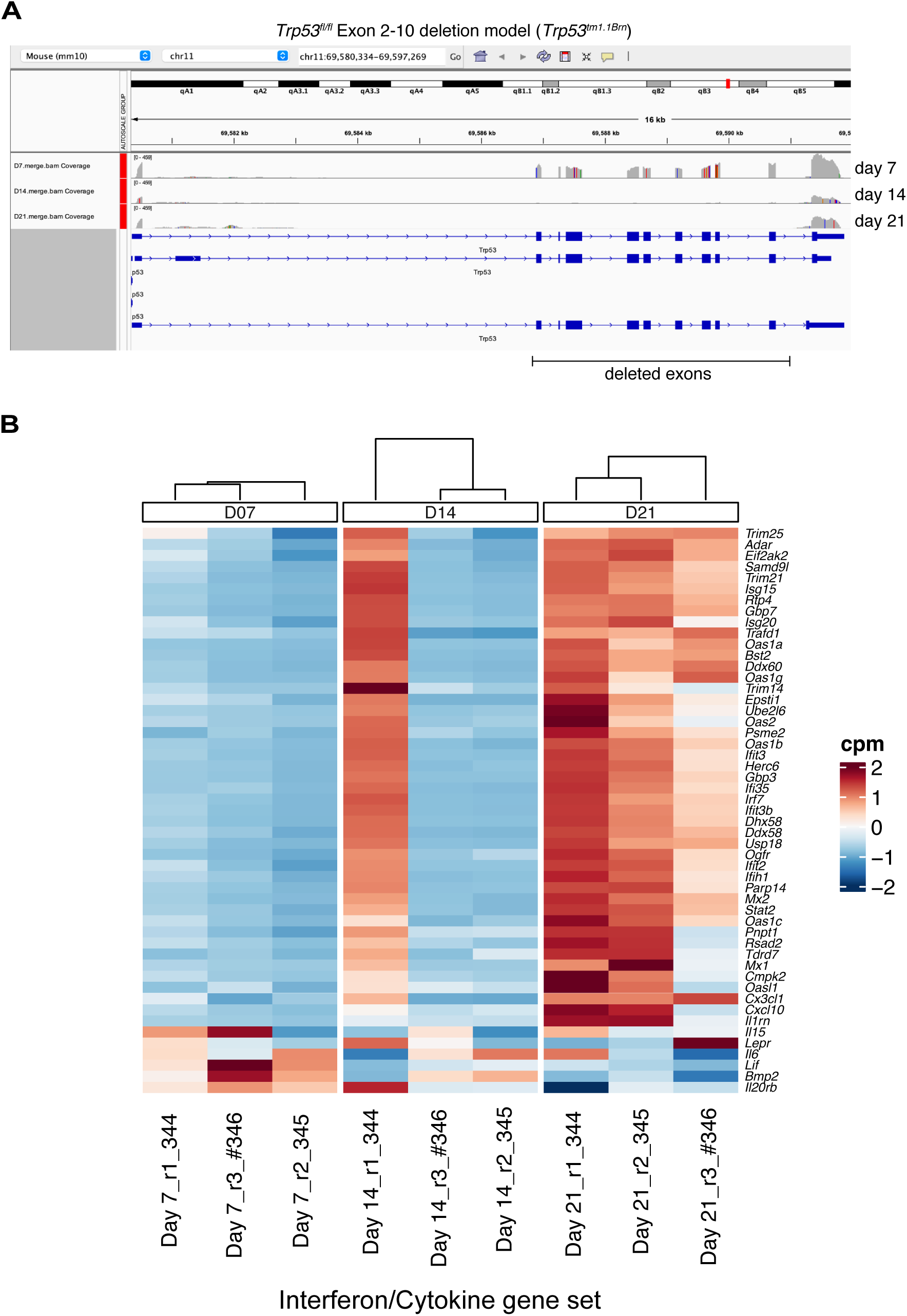
Loss of *Trp53* in osteoblasts leads to an activation of the interferon stimulate gene expression program. **(A)** IGV screen shots of the *Trp53* (p53) locus at day 7, 14 and 21 of tamoxifen treatment. The region deleted by the *Trp53^fl/fl^* allele is indicated. The data show the read counts from RNA-seq. **(B)** The expression of the interferon stimulated gene set in each replicate at day 7, 14 and 21. Data expressed as counts per million.

**Supplementary Figure 7.**
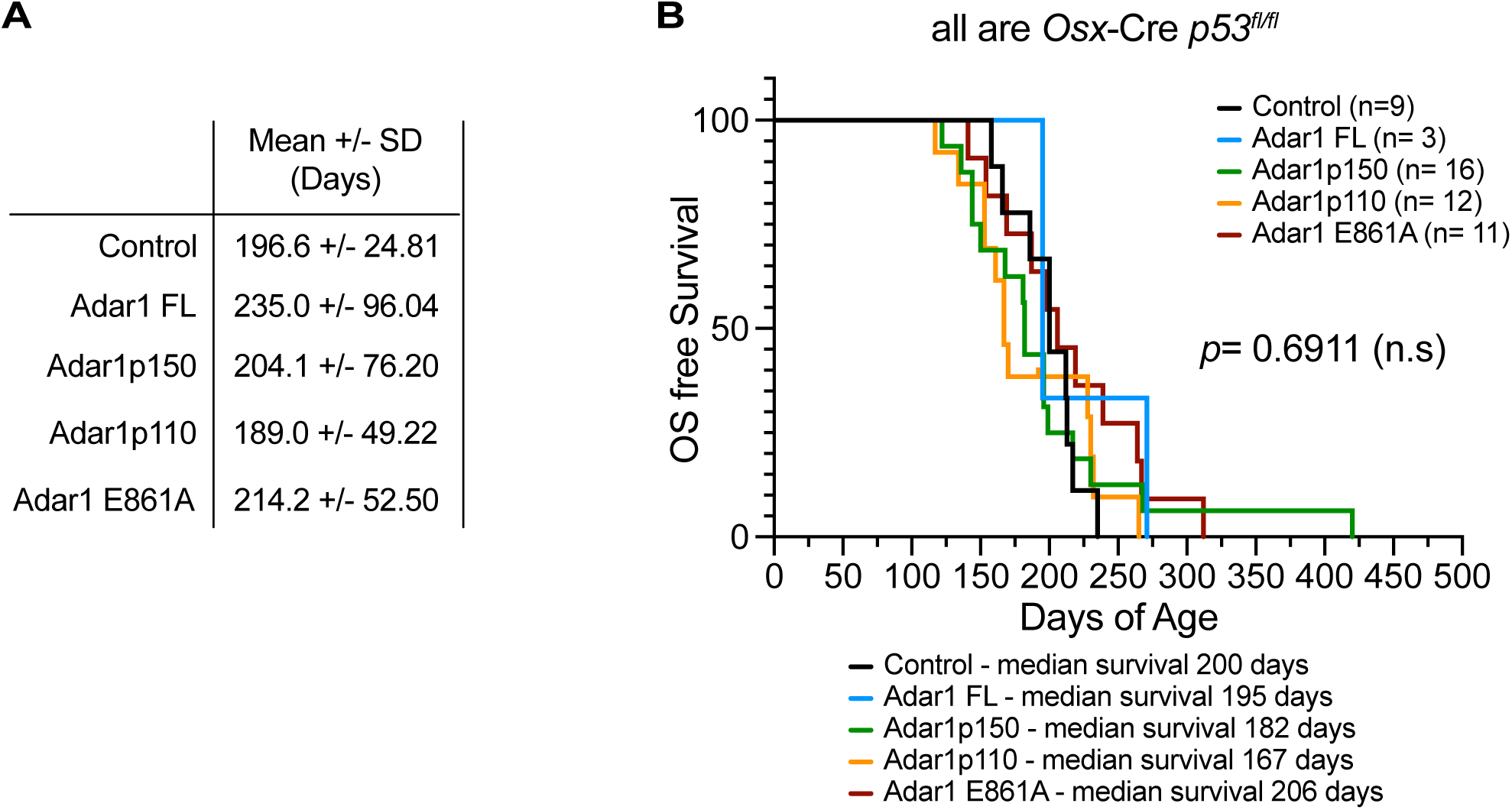
Overexpression of ADAR1 does not accelerate osteosarcoma formation. **(A)** Mean survival +/- standard deviation of each genotype in the KM plot in Figure 7B. **(B)** Kaplan Meier plot based on *Osx1*-Cre *p53^fl/fl^* genotype + the indicated *R26*-Adar1 allele. These can have either *pRb^fl/+^* or *pRb^fl/fl^* genotype. Number as indicated in the inset, no statistical difference. Median survival as determined by Prism KM analysis.

**Supplementary Figure 8.**
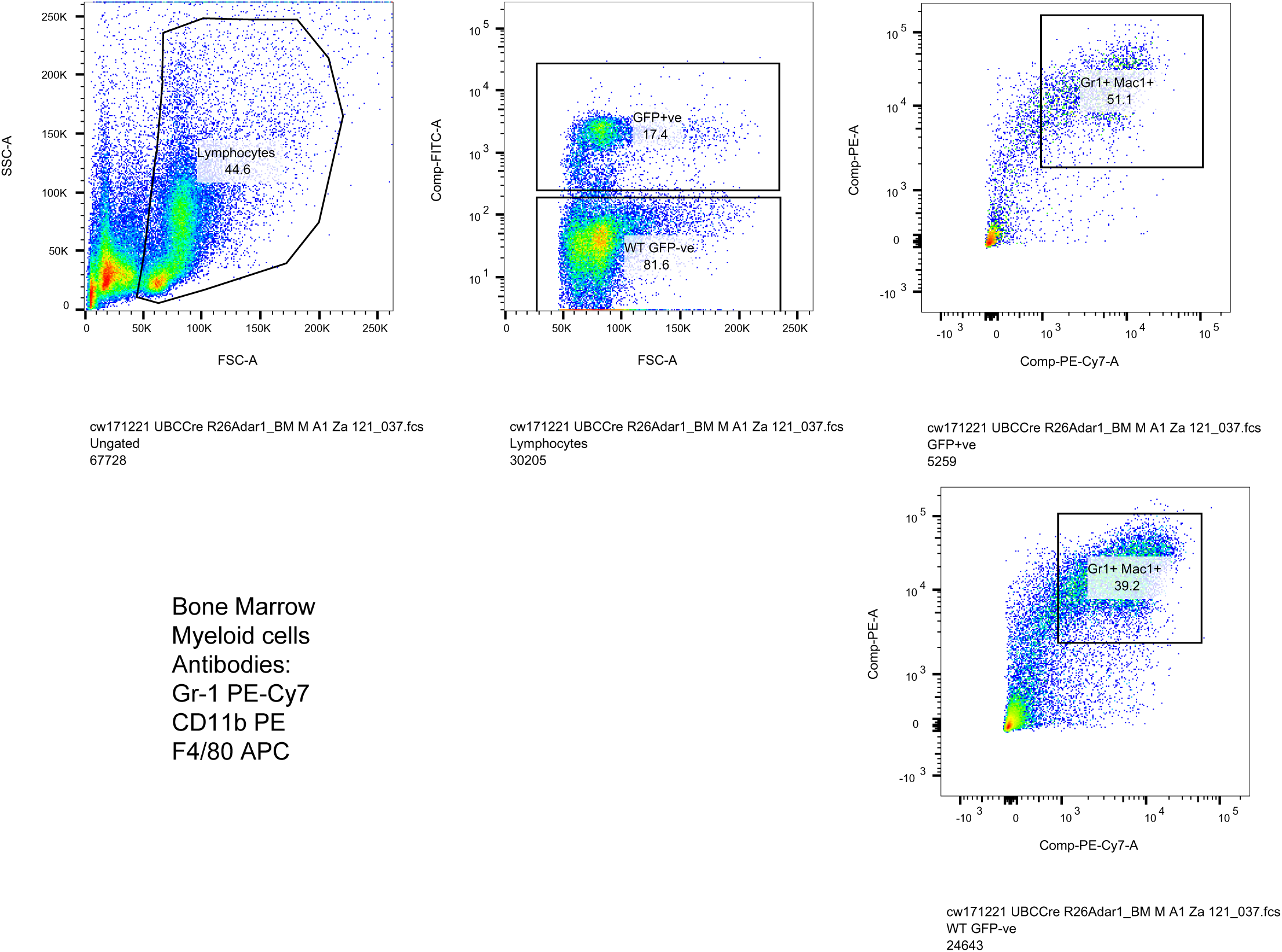
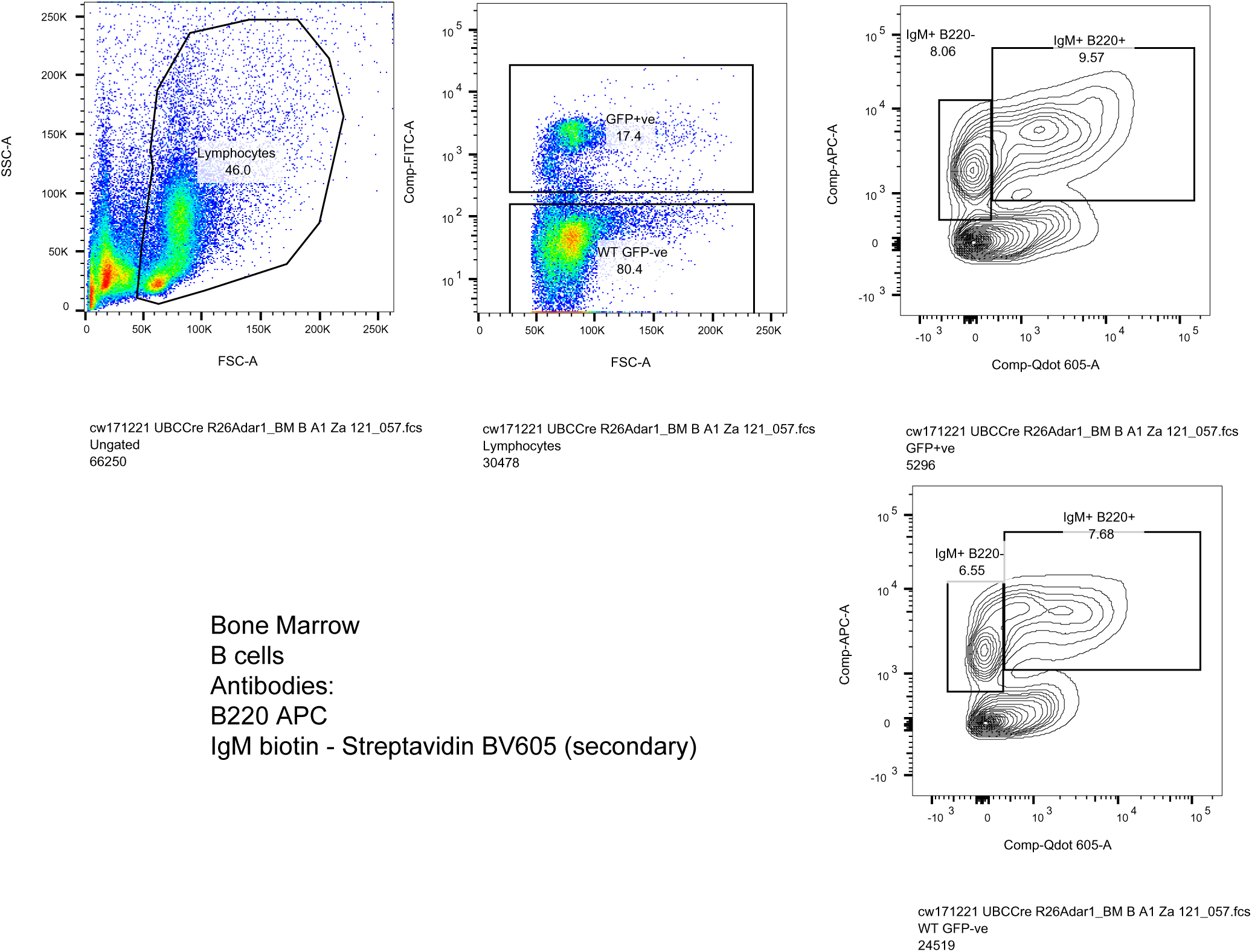
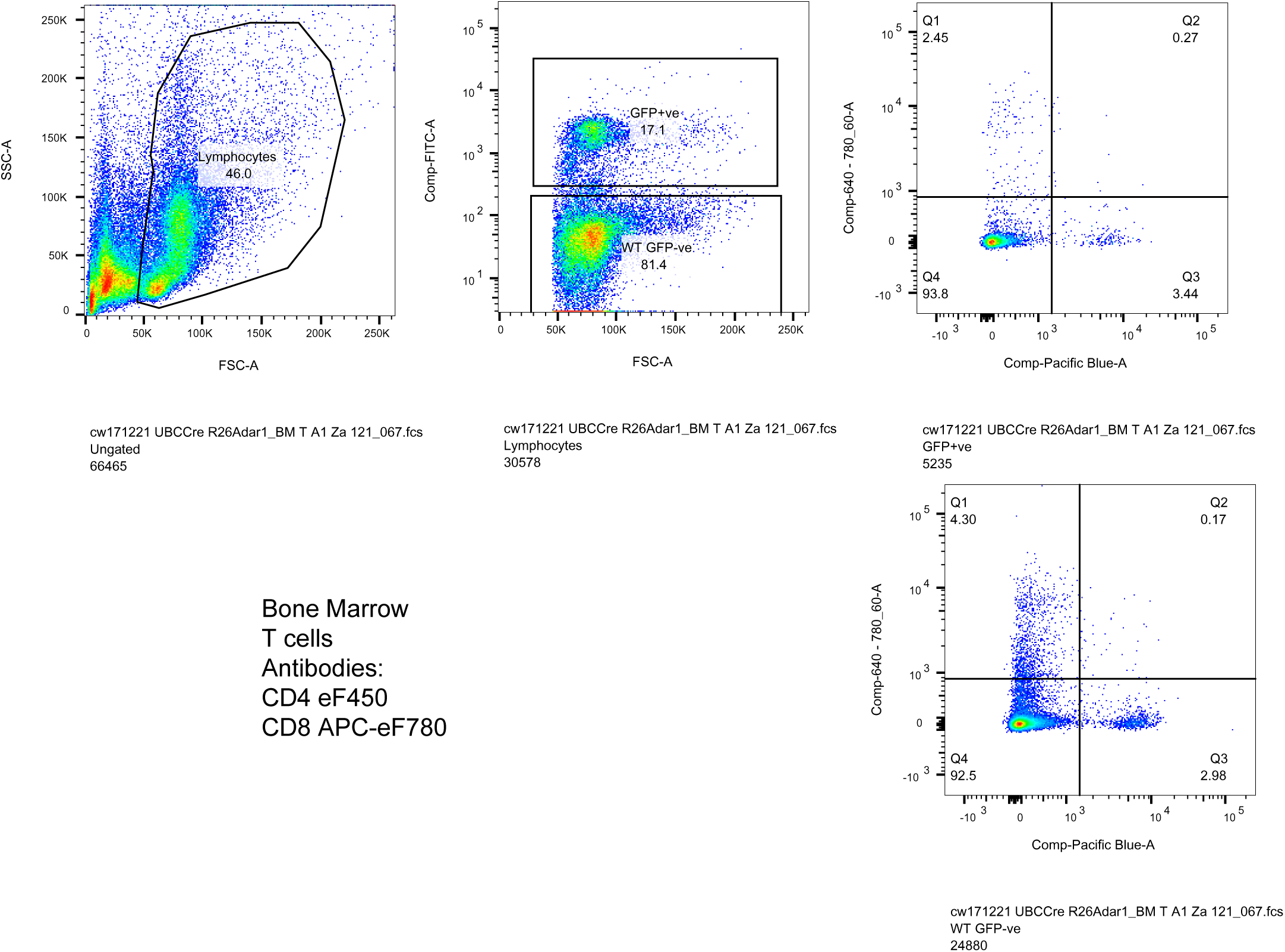
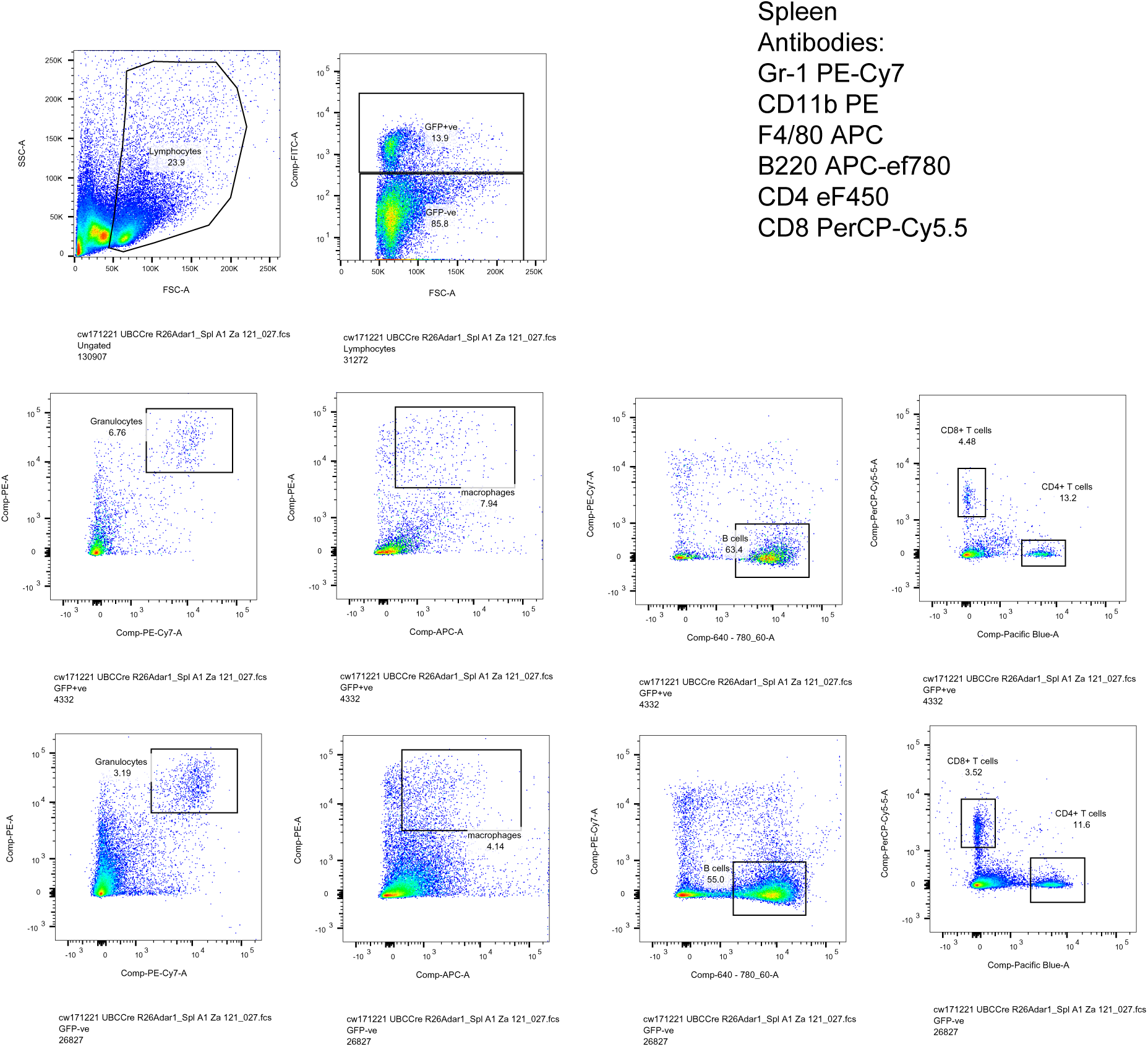
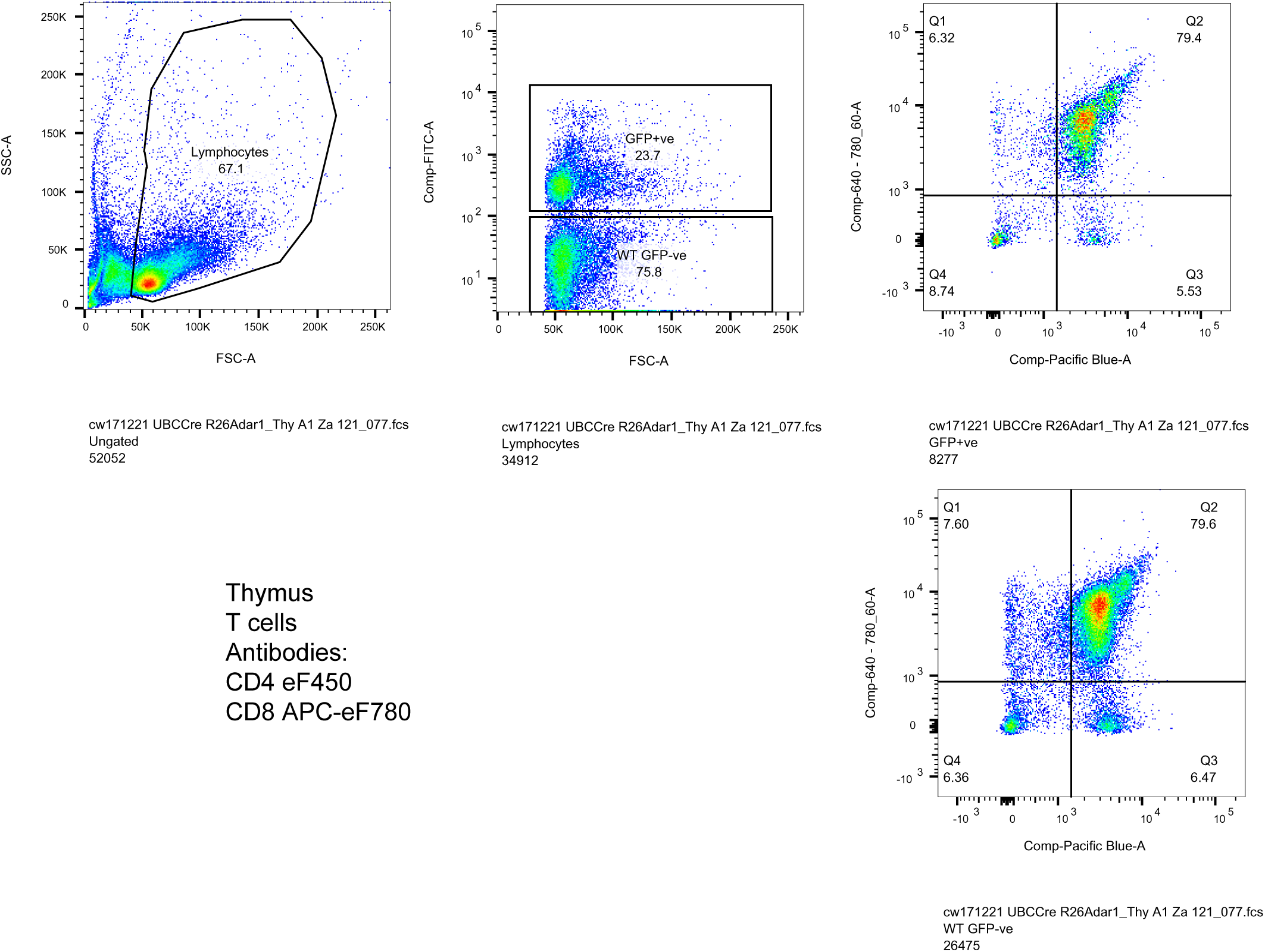
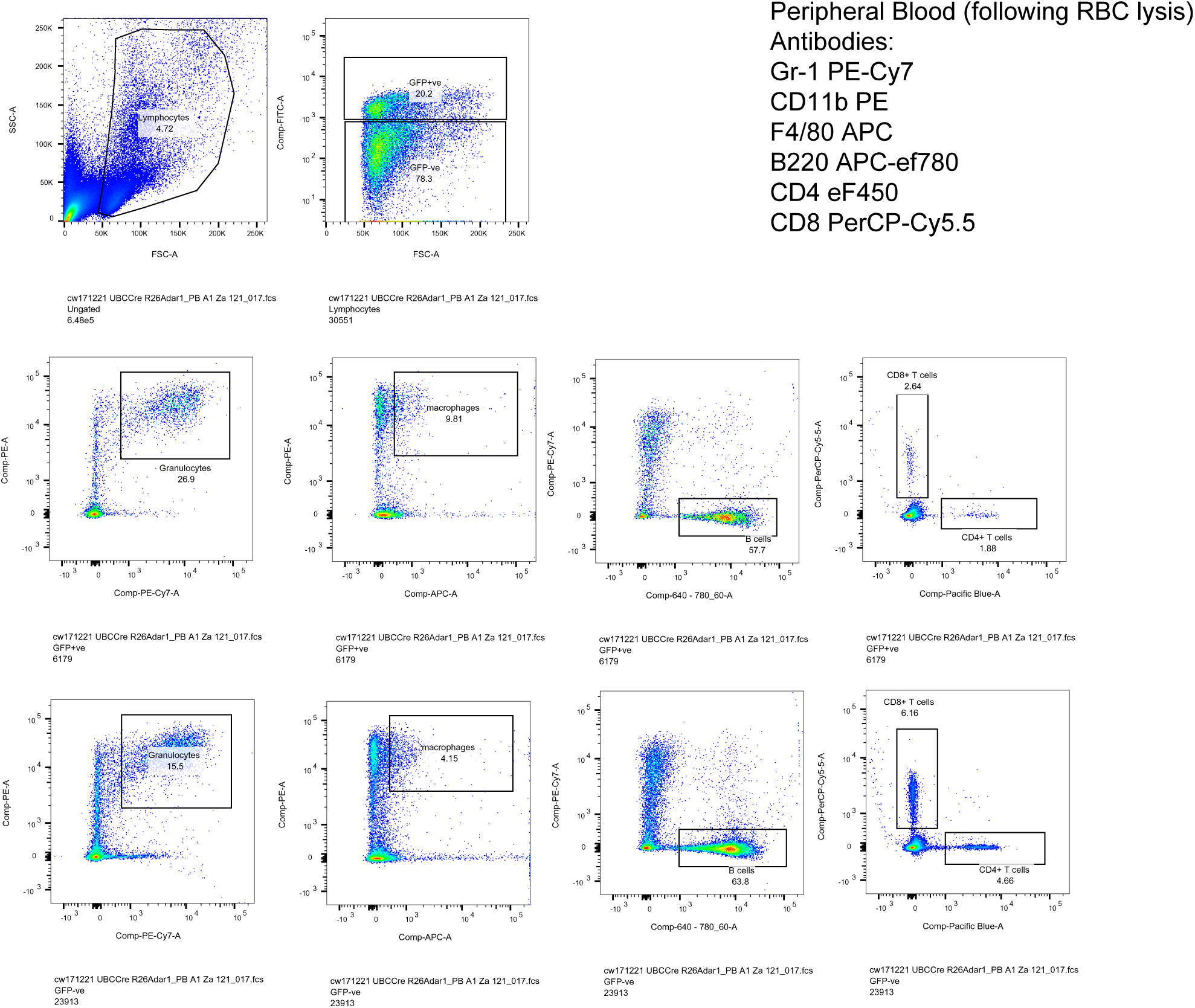
Representative flow cytometry profiles and gating strategies for FACS analysis. Each page is labelled with the organ assessed and the antibodies and conjugates used.

### Tables

**Supplementary Table 1.** Genotyping primers used in this study.

**Supplementary Table 2.** Flow cytometry antibodies, dye conjugates, antibody clone and source used in this study.

### Datasets

**Supplementary Dataset 1:** Gene expression analysis of day 14 liver samples.

**Supplementary Dataset 2:** Editing analysis of day 14 liver samples.

**Supplementary Dataset 3:** Gene expression of osteoblasts.

**Supplementary Dataset 4:** Editing analysis of osteoblasts.

## ACKNOWLEDGEMENT

The authors would like to thank JB Li, L Purton and J Heierhorst for discussion and comment; E Tonkin for technical assistance; P Humbert (La Trobe University) for the *UBC*-CreER^T2^ mouse line; A Kueh at The Melbourne Advanced Genome Editing Center (MAGEC) at the Walter and Eliza Hall Institute as a node of Phenomics Australia for generation of the *R26*-LSL-*Adar1* mouse lines; Monash Antibody Technology Facility (MATF) for purification of ADAR1 antibody from hybridomas; and St. Vincent’s Hospital BioResource’s Centre for care of experimental animals. We thank R Kuehn/Addgene for plasmid distribution. Schematic figures were made using BioRender.com.

## Author Contribution Statement

C.R.W conceptualized the study. S.M.R, J.H-F and C.R.W designed the experiments. S.M.R, A.C., A.G, J.H-F and C.R.W performed the experiments. J.H-F and C.R.W wrote the original manuscript, and all authors reviewed and edited the manuscript; J.H-F and C.R.W were responsible for funding acquisition. J.H.F and C.R.W provided supervision.

